# From species-wide genomics to cellular diversity: characterizing multidrug-resistant extraintestinal pathogenic *Escherichia coli* ST744

**DOI:** 10.64898/2026.11.13.766372

**Authors:** Kristina Nesporova, Bart Steemans, Jana Palkovicova, Kristina Krutekova, Alix Meunier, Adam Valcek, Monika Dolejska, Sander K. Govers

**Author notes:** **co-corresponding authors Corresponding authors and email address** Monika Dolejska, Sander K. Govers.

## Abstract

The species *Escherichia coli* represents a wide and heterogeneous range of strains. Here, we exploited this genomic diversity to examine the distribution of virulence-associated genes (VAGs) and antibiotic resistance genes (ARGs) within a collection of 131,610 *E. coli* genomes. Across lineages of extraintestinal pathogenic *E. coli* (ExPEC), we identified a negative correlation between VAG and ARG content, with ST744 carrying the most resistance genes on average. Phylogenomic analysis of a global ST744 cohort revealed the existence of two main clades, and substantial variation in accessory gene content, resistance-region organization, and plasmid carriage. Subsequent phenotypic characterization revealed that the interpretability of genomic information depends on the trait being considered. Whereas antibiotic susceptibility measurements generally agreed with genomic predictions, limited correspondence was observed between the presence of specific metabolic and stress-associated genes and enhanced growth under such conditions. Image-based profiling also revealed extensive diversity in cell morphology, nucleoid organization and cell-cycle timing across strains, even among nearly clonal isolates. Together, our findings provide an in-depth characterization of *E. coli* ST744, within the broader diversity of *E. coli* and EXPEC, as well as a framework for examining how genomic differences relate to phenotypic diversity within other ExPEC lineages.

## Introduction

*Escherichia coli* displays extensive intra-species diversity, with strains ranging from laboratory workhorses to opportunistic pathogens^1^. The latter category can be broadly subdivided into two main pathogenic groups: intestinal pathogenic *E. coli* (IPEC), causing infections in the gut, and extraintestinal pathogenic *E. coli* (ExPEC), causing infections in other body compartments^1,2^. ExPEC represent the principal cause of urinary tract infections (UTIs), sepsis, and other extraintestinal infections in humans and animals and they are considered a major global health and socioeconomic burden^1,3–5^. In contrast to IPEC, for which the majority of pathovars are identified and classified based on specific virulence-associated genes (VAGs), defining ExPEC strains based on their VAGs remains unreliable^1,3^. Examination of ExPEC distribution across phylogenetic groups (PG) also reveals a heterogeneous pattern, with the majority residing in PG B2, but also many strains belonging to other PGs^4,6,7^. Instead, in the context of ExPEC, it is common to apply sequence type (ST) sub-classification^8^, as only a limited number of STs appear to cause the majority of ExPEC infections (from more than 19,000 existing *Escherichia/Shigella* STs in total)^4^. However, due to their heterogeneous nature, ExPEC strains can hardly be defined as a uniform group, especially at the genomic level^1^.

In general, three main ExPEC groups that capture the main trajectories of ExPEC diversification over time, in terms of PG, VAG content, and antibiotic resistance genes (ARGs), can be identified^4,9,10^:

### Classical high-virulence ExPEC

This group, which has historically been dominant before 2000, contains lineages that have a high, and sometimes specific, VAG content and are typically antibiotic-susceptible^10^. Most lineages are located within PG B2, including ST12, ST73, ST95, ST127, ST141, and are characterized by large, stable accessory genomes rich in adhesins, toxins, iron-acquisition systems, and multiple pathogenicity islands^11,12^. They display strong urovirulence, high adaptability to the human urinary tract, and limited antimicrobial resistance (AMR)^13^. Their (pathogenic) success appears to rely on host specialization and virulence intensity, including efficient colonization, intracellular persistence, and immune evasion, rather than on acquiring ARGs^12,13^.

### Emerging multidrug-resistant (MDR) *ExPEC*

Since the early 2000s, a new wave of ExPEC, (mainly associated with PGs B2 and D, and to a lesser extent with PGs F and B1, including ST38, ST69, ST131, ST405, ST648, and ST1193) have become globally dominant in hospitals and communities^10,14^. These MDR lineages evolved by the stepwise acquisition of ARGs, especially those encoding resistance to higher generations of beta-lactams, typically on epidemic plasmids of IncF or IncX families combined with chromosomal mutations conferring fluoroquinolone resistance^4,10^. Unlike *Classical high-virulence ExPEC*, they often exhibit reduced VAG repertoires, yet display, together with their rich AMR cargo, enhanced metabolic efficiency and plasmid acquisition, enabling long-term persistence under antibiotic pressure^4,15^. They are often referred to as pandemic lineages, as they have been isolated across the globe and can be found in various sources, yet many of them are adapted to the human host. The most notorious example of this group is ST131, which is responsible for a significant proportion of ExPEC infections^4,16^, is known to be atypical in comparison to the other members of PG B2 (lower VAG cargo and a typical MDR profile)^17,18^, and is commonly subdivided into specific human-associated clades (e.g., ST131-C1 or ST131-C2)^16,19^.

### One-Health high-risk ExPEC

commonly also referred to as international high-risk clones. This most recent and still expanding group is linked to non-B2/D PGs (mainly A and C, but also F and B1), with ST167 and ST410 being the best-described representatives so far^10^. These lineages blur the boundaries between commensal, environmental, and pathogenic *E. coli*^20^, and are frequently encountered in wild and food-producing animals and environmental niches, while being increasingly recovered from human infections^21,22^. The high-risk lineages are commonly linked with extensive antibiotic resistance on diverse plasmids, including resistance to last-resort antibiotics such as carbapenems^10,23^. In some cases, their AMR cargo even surpasses that of emerging MDR lineages. Their VAG content is often limited, but they display broad ecological flexibility, stress tolerance, and rapid horizontal gene transfer, often acquiring partial virulence modules and metabolic innovations that support cross-host survival^10,24,25^.

Across the above-described groups, ExPEC lineages appear to balance virulence and resistance as alternative adaptive strategies^26,27^. Yet, this remains mostly based on anecdotal evidence coming from a small subset of ExPEC strains or lineages. Therefore, in this study, we performed a large-scale, species-wide genomic analysis to assess the interplay between virulence and resistance. This analysis revealed a striking general negative relationship between the VAG content and the number of ARGs across ExPEC STs. We subsequently focused on ST744, which showed the highest average ARG content and is also repeatedly detected globally across human, animal and environmental reservoirs, and associated with clinically important AMR determinants^28,29^. However, despite all this, it has remained a lesser studied *One-health ExPEC*, and the global population structure, genomic characteristics, and phenotypic properties of ST744 had remained poorly resolved. All of this is addressed below.

## Results

### Species-wide analysis of VAG and ARG content in *E. coli*

To explore the genomic diversity of *E. coli* in terms of VAG and ARG content, we used our recently assembled and annotated large-scale genome collection consisting of 131,610 *E. coli* genomes^7^. This collection contains a broad range of genomes, with many annotated as *IPEC* (42,791 genomes) or *ExPEC* (20,523 genomes), but also genomes of which the clinical status could not reliably be determined (*ND*, 65,828 genomes) and several other minor groups. In a first step, we evaluated the VAG content of the genomes by performing a principal component analysis (PCA) using the presence or absence of all potential VAGs as input (Fig. 1A). The resulting plot adequately captured different phylogroups (Fig. 1A, left panel), but also revealed a clear differentiation between *IPEC* and *ExPEC* (Fig. 1A, right panel), with many *ExPEC* genomes clustering among other non-IPEC genomes, which is in line with our previous findings that focused solely on genomes containing a type VI secretion system^7^. In a subsequent step, we examined the distribution of the number of VAGs and ARGs, within all genomes of our collection (Fig. S1A), and within *IPEC* and *ExPEC* genomes specifically (Fig. S1B-C). This analysis revealed that *IPEC* genomes tend to have a higher VAG content and rarely carry any ARGs (Fig. S1B). In contrast, *ExPEC* genomes displayed a more dispersed pattern. While the majority of ExPEC genomes was located in areas with limited ARGs and elevated VAGs (though not as high as *IPEC*), we also observed many genomes with moderate VAG content and a high number of ARGs (Fig. S1C). Moreover, the ARG distribution of *ExPEC* genomes displayed a bimodal pattern, meaning that if ARGs were detected, these genomes typically contained higher numbers of ARGs (more than the average of 3-4 ARGs across all *ExPEC* genomes).

**Figure 1.**
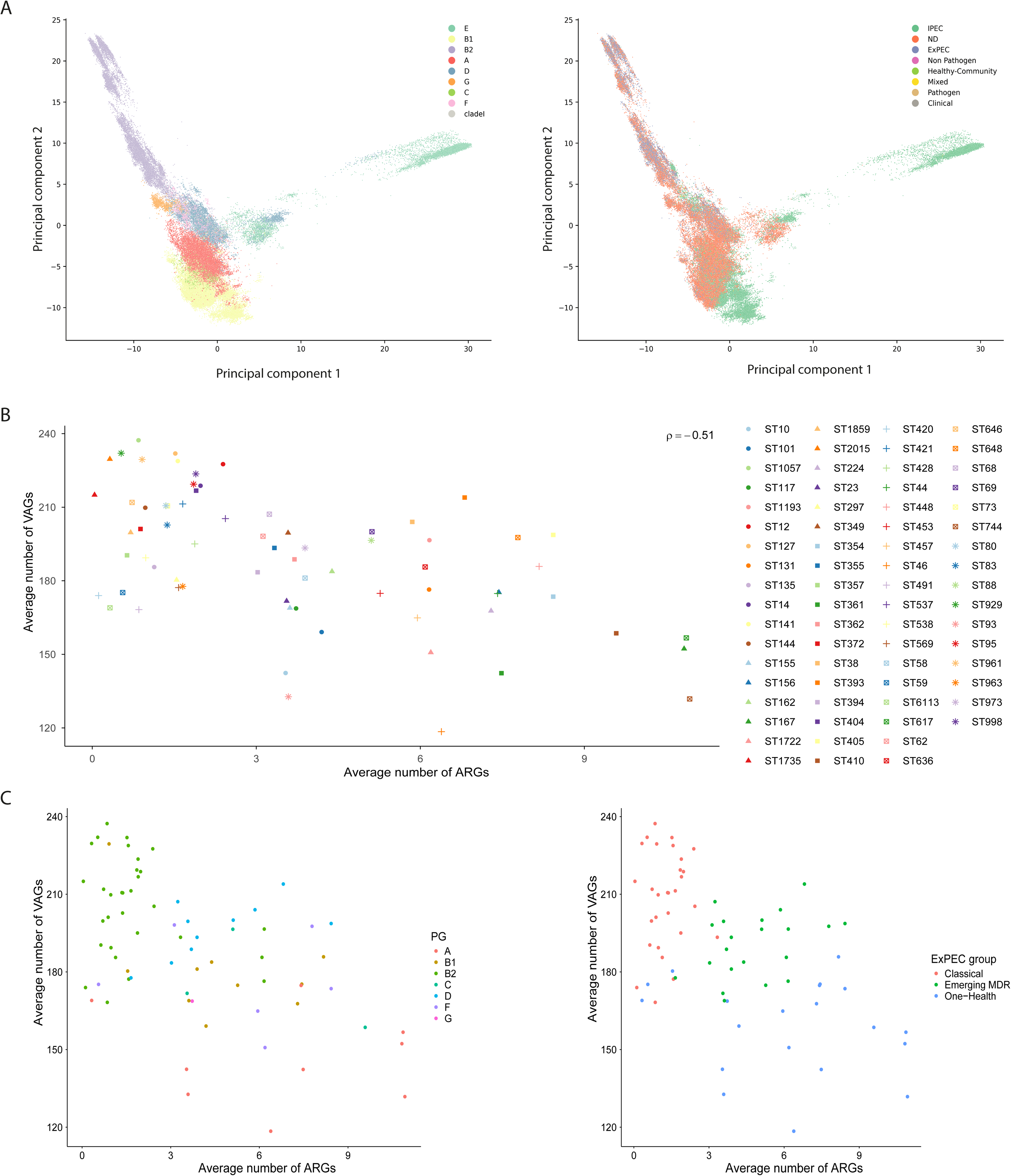
Virulence and resistance cargo-based analysis across *E. coli* population and dominant ExPEC STs. **A)** Scatterplot displaying the PCA decomposition for phylogenetic group (left) and Source-curated (right) based on 695 VAGs identified using the VFDB *E. coli* database. **B)** Relationship between the average number of detected virulence-associated genes (VAGs) and antibiotic resistance genes (ARGs) for the 70 most dominant ExPEC sequence types, the Spearman correlation is indicated in the right top corner. **C)** Relationship between the average number of detected virulence-associated genes (VAGs) and antibiotic resistance genes (ARGs) for the 70 most dominant ExPEC sequence types, highlighting the phylogenetic groups (left) and the three ExPEC groups (right).

### Apparent trade-off between VAG and ARG content in ExPEC

Previous work indicated that only a small fraction of STs is responsible for the majority of ExPEC infections (20 STs linked to 85% of ExPEC genomes^4^). We used our own large-scale collection of 20,523 *ExPEC* genomes and observed a similar trend, as the first 8 STs (ST131, ST73, ST69, ST95, ST1193, ST127, ST38 and ST12) already represented 50.3% of *ExPEC* genomes (from a total of 1,666 STs for which at least one genome was labelled as *ExPEC*). To compare directly, our 20 most common ExPEC STs covered 67.2% of *ExPEC* genomes, while 98 and 222 STs would be needed to reach >85% and >90% of *ExPEC* genomes, respectively (Fig. S1D).

Based on this analysis, we focused on the 70 most common ExPEC STs that had minimally 35 genomes labelled as *ExPEC* to ensure a reasonable representation. Together, these 70 STs covered >82% of *ExPEC* genomes. Plotting the average number of ARGs versus the average number of VAGs for these 70 most dominant ExPEC STs revealed a negative correlation between them (Fig. 1B). This relationship suggests that, on average, STs with more ARGs tend to have less VAGs, and vice versa (Fig. 1B). Furthermore, the distribution of STs and PGs matched that of previously described patterns with a division of ExPEC into three main groups. Highly virulent STs were almost exclusively from PG B2 and represent the *Classical high-virulence ExPEC* with a high number of VAGs that dominated the early ExPEC era (Fig. 1C). STs with attenuated virulence cargo and a typical MDR profile come from different PGs (B1, B2, F, C, and D) and represent the *Emerging MDR ExPEC* (Fig. 1C). This group contains three atypical PG B2 STs (i.e., ST131, ST1193, and ST636), which deviate from other members of the same PG in terms of ARGs and display a more moderate VAG content. Both ST131 and ST1193 are successful global ExPEC lineages, with ST131 being described as the most successful MDR lineage of all time and ST1193 as an emerging lineage following in its footsteps^4,16,30,31^. A final group of STs, which represent the *One-Health high-risk ExPEC*, exhibited lower average numbers of VAGs but increased and sometimes even extreme levels of ARGs (Fig. 1C). These *One-Health* lineages mostly belong to PG A, but also PG B1, C, and F. Overall, ST744 exhibited the highest number of ARGs. While this ST has been previously reported as a high-risk lineage^32^, it has not yet been subjected to a detailed global examination.

### Phylogenetic analysis of a ST744 global cohort reveals two main clades

Given its remarkable number of ARGs, we set out to further examine ST744, and characterize the genomic and phenotypic properties of its members. For this, we first performed a phylogenetic analysis of a global cohort of 892 ST744 genomes (Fig. 2; Table S1). This phylogenetic analysis was based on 2,135 core genes present in all 892 strains from a total of 26.200 predicted genes (Table S2). The resulting phylogenetic tree revealed the existence of two main clades, which cluster according to two serotypes, O101:H9 and O101:H10 (Fig. 2), with H9 being detected in 55.8% of strains, H10 in 40.6% and O101 in 96.6%. The third most common serotype, O101:H25, was represented by a clade of only 20 strains (Fig. 2). Within the strains with a O101:H9 serotype (Fig. 2, green), a small yet distinct clade of 8 phylogenetically distant strains can be observed (Fig. 2, darker green). The SNP difference between this small distant O101:H9 clade and all other genomes ranges around 3.000, with a (global) maximum of 3983 SNPs between FB4014AA and IB9790AA (Table S3), a genome from a more distant subclade of O101:H10. These distant strains were represented by isolates from different sources and countries, with their common denominator being the absence of chromosomal mutations causing resistance to fluoroquinolones and fosfomycin (further details presented below). This clade likely represents the evolutionary older genomes of ST744, as the switch from fluoroquinolones-susceptible to fluoroquinolones-resistant was shown to be an important evolutionary milestone linked with clade separation for other ExPEC lineages, including ST131^16,18,33^ and ST410^10^.

**Figure 2.**
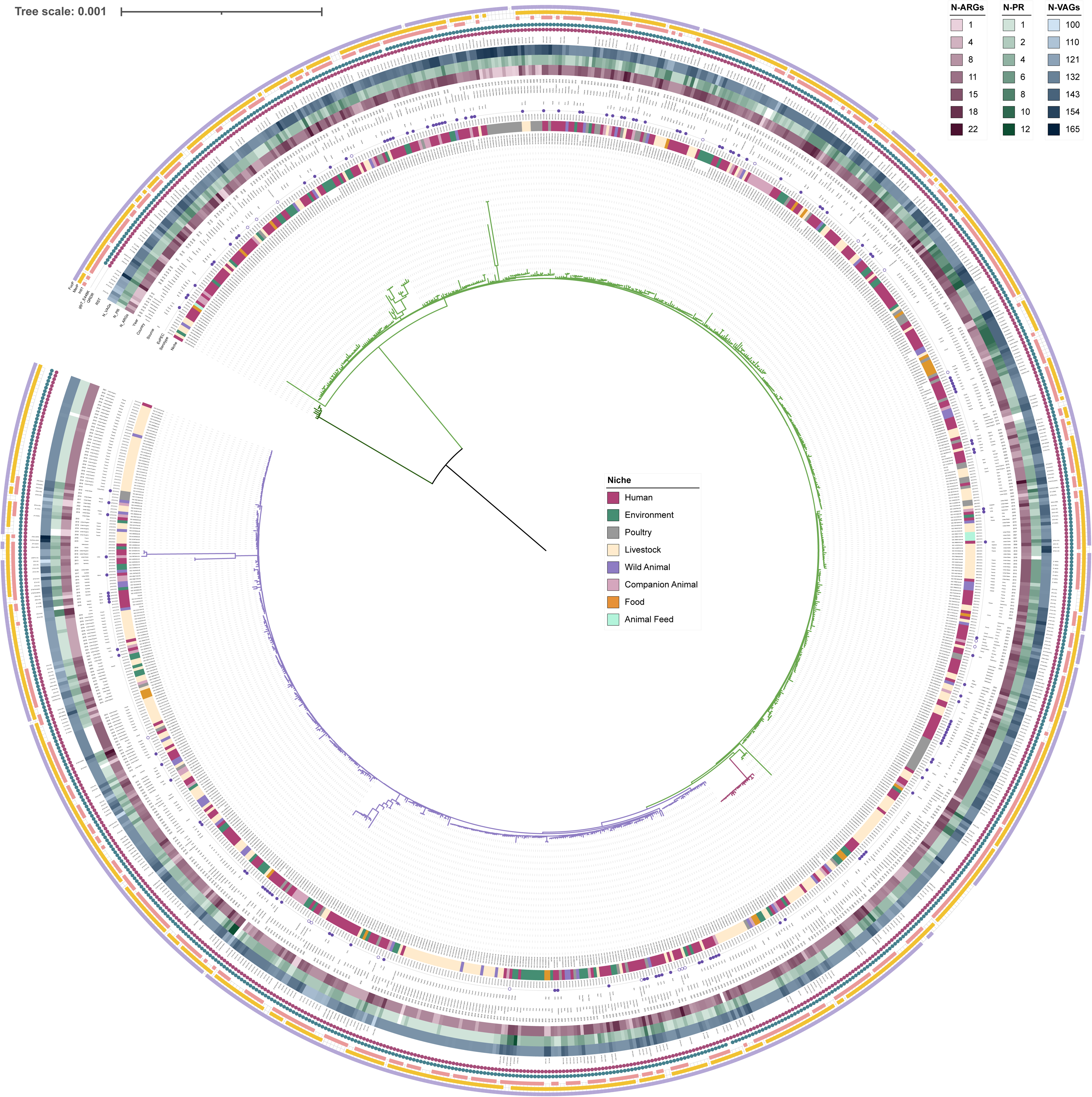
Phylogenetic analysis of ST744 collection. The branches are coloured according to the dominant serotype: O101:H9 (green, a distinct clade of this serotype in dark green), O101:H10 (violet) and O101:H25 (pink). The metadata in order from the most inner circle resents Niche, Serotype, *ExPEC* status (ExPEC, with full violet circles representing *ExPEC*-labelled strains, while empty violet circles represent additional clinically relevant strains without further specification), Source of isolation (Source), Country of origin (Country) and Year of isolation (Year). Number of detected ARGs (N_ARGs), plasmid replicons (N_PR) and VAGs (N_VAGs) are highlighted in the form of a heatmap. Further, RST of F plasmids is specified and the presence of specific markers is shown as full circle/square including QRDR point mutations (*gyrA*D87N, *gyrA*S83L, *parC*S80I, *parC*A56T, pink circle), and a point mutation causing resistance to fosfomycin (*glpT*_E448K, turquoise circle), presence of *intI1* integrase (IntI1, orange square), the presence of the representative gene *merP* from the mercury resistance operon (MerP, yellow square) and the presence of the specific *fucP* transporter (FucP, violet square).

The SNP difference within the two main clades typically ranged from tens to hundreds of SNPs (Table S3). Within clades, we also identified numerous clonal pairs among the genomes (0-10 SNPs), typically for genomes coming from the same *Niche* but in some case also for genomes with different *Niches*. This includes approximately 15 clades where clonal pairs from different *Niche* came from the same *Country* and *Year* suggesting a possibility of an epidemiological link (e.g., FC1258AA *Human* and FC1522AA *Companion animal* in Portugal, FC0999AA *Human* and EC9529AA *Environment* in Ecuador, AC0164AA *Wild animal* and AC0210AA *Livestock i*n UK; Table S4). Notably, for the UK, clades with strains from 2018 from *Livestock* or *Wild animals* were clonal to *Human* strains from 2021 (Table S4). A similar clonal pair from Spain came from *Poultry* (WB3421AA, 2015) and *Human* (GC2550AA, 2021) while the strains even share the same F18:A6:B1 plasmid (Table S4). To further probe genomic properties that are associated with one of the two main clades, we performed a genome-wide association study (GWAS). This analysis revealed that genes strongly associated with the H10 serotype (Table S5) are predicted to encode a set of DNA-cleaving enzymes linked to restriction-modification systems or defense against mobile genetic elements (predicted proteins XOG39217.1, XOG39216.1, UBO44270.1, Table S5), or are mobile genetic elements themselves, in the form of IS*200*/IS*605* family transposase (XOG39187.1). Genes strongly associated with H9 serotype (Table S6) were linked to presumable functions of adhesion (*yeeJ*), quorum sensing (*lsrB*), membrane transport (*yeeO*), carbohydrate metabolism (D-inositol-3-phosphate glycosyltransferase *mshA*) and mobile genetic elements (IS*Kpn42*).

### Correspondence between presence of antimicrobial resistance genes and resistance phenotypes of ST744

ST744 is a lineage with a high level of AMR, with 889 genomes (99.7%) being classified as MDR (detection of ARGs/point mutations conferring resistance to 3 or more groups of antibiotics). Predicted resistances to fluoroquinolones, fosfomycin, beta-lactams, aminoglycosides, sulphonamides and tetracyclines reach over 90% prevalence (Fig. S2, Table S1). On average, 11 ARGs were detected in our dataset of 892 genomes, with a maximum of 22 ARGs in two strains (Fig. 2, Table S1). Only two strains did not contain any ARGs, yet they both carry point mutations providing resistance to fluoroquinolones and fosfomycin. In fact, nearly all genomes (881; 98.8%) carried a point mutation, *glpT*E448K, which confers resistance to fosfomycin (Fig. 2, Table S1). Point mutations linked to fluoroquinolone resistance in the quinolone resistance determining regions (QRDR) were also common, with 884 genomes (99.1%) carrying the same four QRDR mutations, *gyrA*D87N, *gyrA*S83L, *parC*S80I, *parC*A56T (Fig. 2, Table S1). The first three of these QRDR mutations are well known to provide clinically relevant levels of fluoroquinolone resistance, even when present individually^10,34^. Several other QRDR mutations were also detected, with some being dispersed across the phylogenetic tree and some being parts of smaller clades. An example of the latter is *parE*S458A, which is linked to a single clade of 24 isolates (Fig. S2). In addition to chromosomal point mutations, plasmid-mediated quinolone-resistance genes occurred, although less frequently and in a dispersed fashion across the phylogenetic tree (Fig. S2). We identified 79 genomes carrying a variant or multiple variants of *qnrA*, *qnrB* or *qnrS.* From these, *qnrS1* was the most dominant (60 genomes).

Among other clinically important ARGs prioritized for monitoring^35^, we found extended-spectrum beta-lactamase (ESBL) and/or AmpC type beta-lactamase (AmpC) genes encoding resistance to 3^rd^ and higher generations of beta-lactams in 442 genomes (49.6%), while 62 genomes (7.0%) carried different variants of carbapenemases, and 113 (12.7%) carried *mcr* genes encoding mobile resistance to polymyxins (Fig. S2), represented mostly by *mcr-1.1* (11.9%) and also by *mcr-1.2*, *mcr-1.32*, *mcr-3.1*, *mcr-3.2* and *mcr-3.5*. For ESBL genes, CTX-M variants were the most common, with *bla*_CTX-M-55_ (134 genomes), *bla*_CTX-M-15_ (73 genomes) and *bla*_CTX-M-1_ (66 genomes) being the most prevalent. Their distribution did not display a clear phylogenetic pattern, as they were found in genomes of both dominant serotypes. The multiple genes encoding different carbapenemases included *bla*_NDM-1_ (18 genomes), *bla*_OXA-48_ (14 genomes), *bla*_NDM-5_ (13 genomes) and several others detected in less than 5 genomes (*bla*_GES-5_, *bla*_IMP-4_, *bla*_KPC-2_, *bla*_KPC-3_,, *bla*_NDM-7_, *bla*_OXA-181_, *bla*_OXA-232_and *bla*_VIM-1_). Similar to the ESBL genes, we found no distinct phylogenetic pattern in the distribution of the carbapenemases. For one of them, *bla*_OXA-48_, we did observe a clear geographical link as 11 of the 14 genomes harboring this gene came from Spain (Fig. S2).

While plasmids play notoriously important roles in the dissemination of various ARGs^36–38^, ST744 genomes carried multiple ARGs in a specific chromosomal region. Besides ARGs, this region was characterized by the presence of a mercury resistance operon (at least 1 gene present in 807 genomes, with the full operon present in 755 genomes; Fig. 2, Fig. 3A, Table S7) and, in the majority of cases (630 genomes), an integrase-encoding *intI1* gene (Fig. 2, Fig. 3A, Table S7). By identifying regions of difference (RODs) between our two long-read sequenced genomes and a common background (MG1655 lab strain), we found that these chromosomal regions are quite variable. In SGL280 (GVB10h), the region comprised approx. 38,630 bp and contained *intI1* and 10 ARGs, while for SGL242 (7727), the ROD consisted of approx. 20,670 bp and contained only five ARGs, of which four were also found in SGL280 (i.e., *aadA5*, *sul1*, *tet*(B) and *dfrA17*). Both RODs shared an approx. 7,620 bp long region that contained the *mer* operon and genes encoding multiple mobile genetic elements (MGEs). The rest of the respective RODs possessed major differences and rearrangements, and could not be aligned. Besides the prevalent mercury resistance markers, we also often detected genes associated with resistance to other heavy metals (e.g., arsenic, antimony, tellurite, copper, nickel, cobalt, and silver), albeit much less prevalently (Table S8). Next to the mercury operon, we found the second highest prevalence (in 11-12% of genomes) for eight genes linked to silver resistance (Table S8).

**Figure 3.**
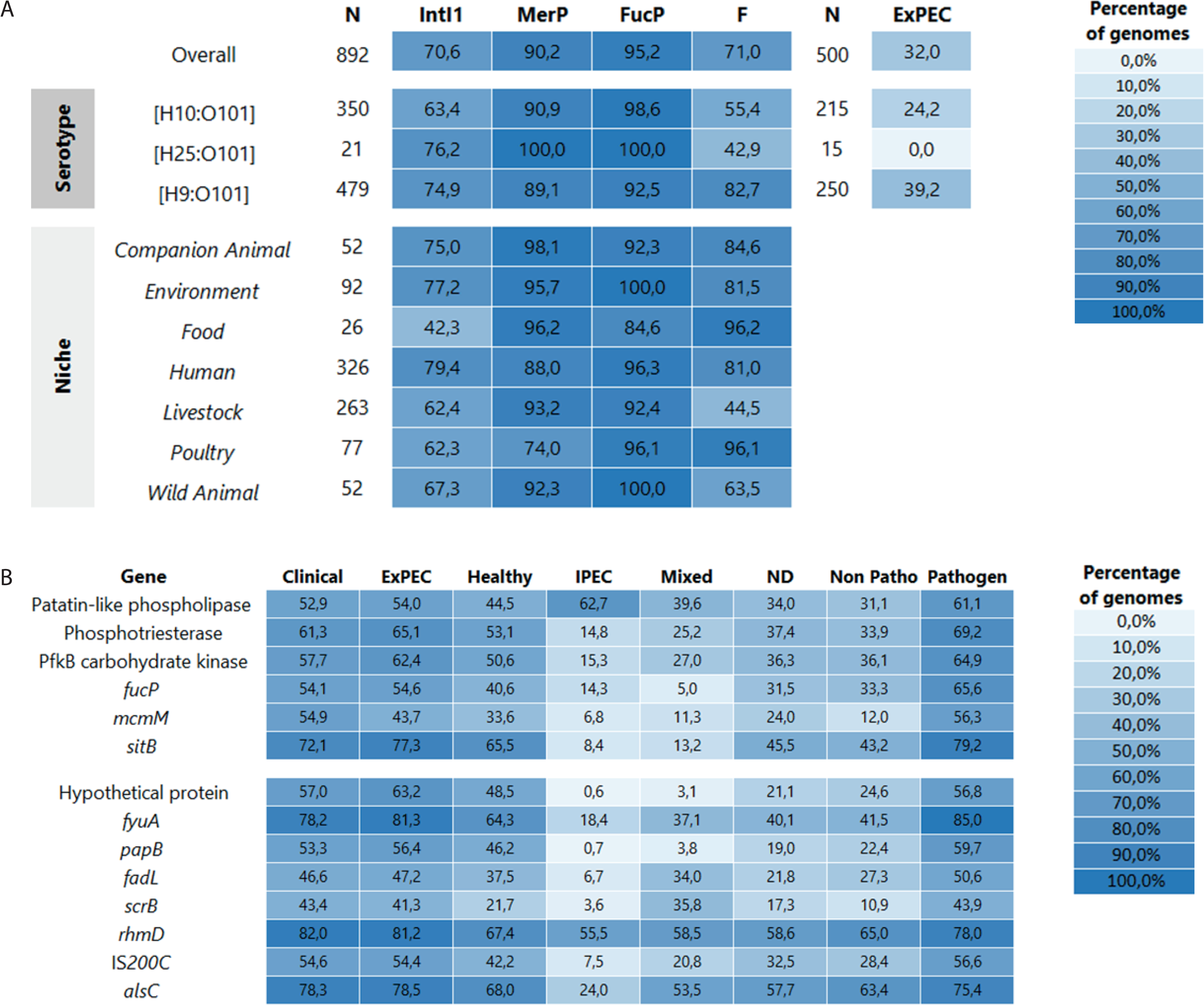
Prevalence of important genomic markers among ST744 and prevalence of *ExPEC*-associated genes. **A)** Prevalence of selected genomic markers across the collection of 892 ST744 genomes (Overall), for the three dominant serotypes (Serotype) and for the different niches (Niche). The first column (N) contains the number of genomes for each category, the following four columns show prevalence in percentage for IntI1, MerP, FucP and F plasmids. The last two columns account only for a subset of genomes where the clinical status could be determined, with N here representing the number of genomes and ExPEC prevalence of *ExPEC*-labeled genomes for the specific groups; the percentages are not calculated for *Niche* category as it was used to assign the *ExPEC* labels. **B)** Prevalence of selected genomic markers found in ST744 and associated with *ExPEC* genomes in our large EB collection (20523 *ExPEC* genomes in total). The prevalence is presented in percentages, the first six rows represent genes identified using a comparative genomic approach and the rest represent selected additional genes identified based on the large-scale screen of the accessory genome.

Given that all antimicrobial resistances so far were only predicted based on the bioinformatic detection of ARGs, we conducted phenotypic testing using commercial minimum inhibitory concentration (MIC) assays for 32 ST744 isolates (Fig. 4, Table S9). The results of these were largely in line with our genomic characterizations. We observed high levels of resistance to fluoroquinolones, with all isolates being resistant to ciprofloxacin. The level of resistance to extended-spectrum penicillins such as ampicillin and piperacillin was also high with 31 isolates being resistant to them, while the single susceptible isolate correspondingly lacked any relevant resistance genes or chromosomal mutations. Similarly, all but one isolate that did not carry any *tet* genes showed MIC to tetracycline greater than 32 mg/L. A total of 23 isolates showed a MIC greater than 4/76 mg/L for the combination of trimethoprim/sulphamethoxazole which was mostly consistent with the presence of *dfrA* and/or *sul* genes variants, despite some examples where a susceptible phenotype was observed even when both genes were present (e.g., SGL242, SGL243) (Table S9). Fosfomycin was not included in the MIC assay. Resistance to colistin was detected in 16 isolates, which corresponded to the presence of *mcr-1* (Fig. 4). Six isolates showed resistance to gentamicin, amikacin, netilmicin and tobramycin. While aminoglycoside resistance genes were detected in almost all of the isolates, they were mostly linked to streptomycin or kanamycin resistance, which were not tested in the MIC assay. A total of 13 isolates showed resistance to cefotaxime (extended-spectrum beta-lactam), with 12 of them carrying a variant of *bla*_CTX-M_ and one carrying a combination of *bla*_CMY-2_ and a carbapenemase *bla*_IMP-4_. Correspondingly, this isolate was also the only one in our set of 32 displaying resistance to carbapenems (meropenem and ertapenem) (Fig. 4).

**Figure 4.**
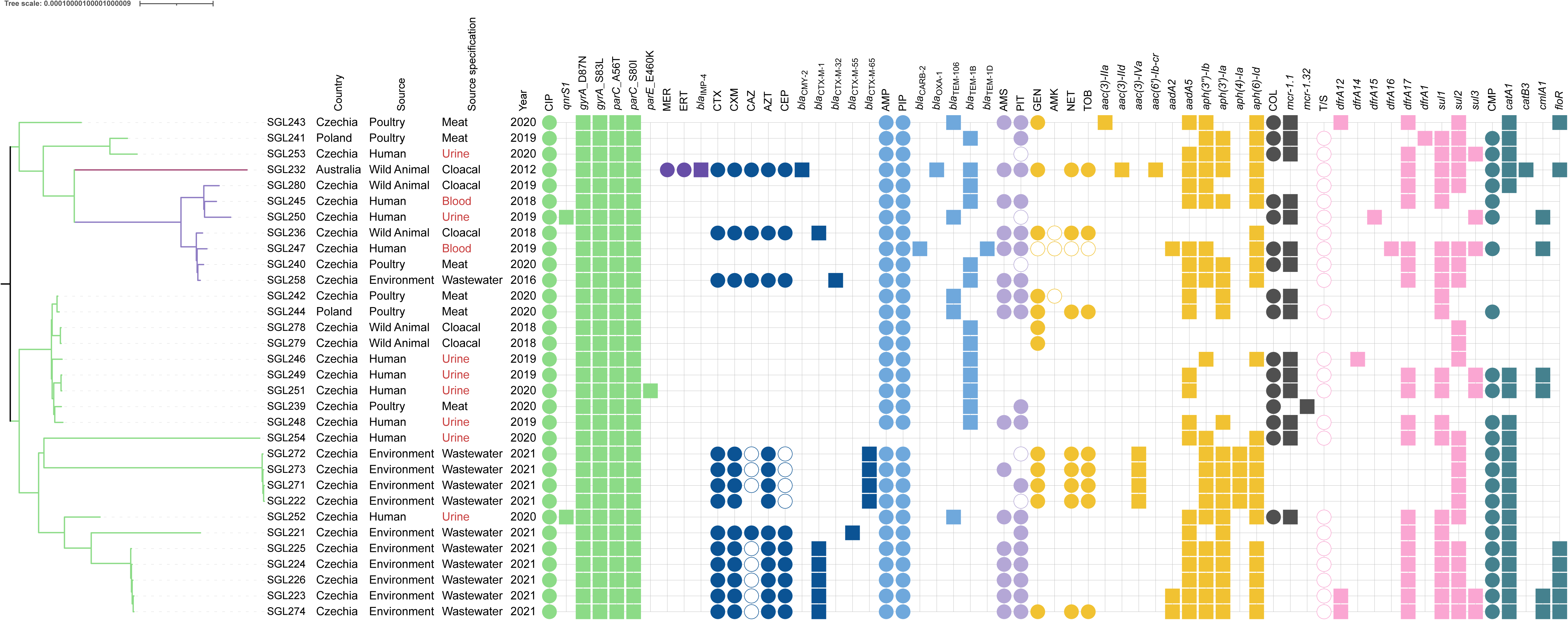
Comparison of antibiotic resistance genomic markers and phenotype. The phylogenetic tree was constructed for 32 strains in our collection and the branch colour highlights the serotype (O101:H9 green, O101:H25 pink, O101:H10 violet). The accompanying metadata contains Country of origin, Source of isolation, Source specification and Year of isolation. The following part depicts whether the strain is phenotypically resistant (full circle), intermediate (empty circle) or susceptible (no circle) to a given antibiotic, for each antibiotic group which is highlighted in the same (or similar for the different groups of beta-lactams) colour, the presence (full square) of relevant genomic markers is highlighted. From left to right, the antibiotic groups include fluoroquinolones (green), represented by ciprofloxacin (CIP); beta-lactams (violet-blue) which comprise of carbapenems (dark violet) represented by meropenem (MER) and ertapenem (ERT), extended-spectrum cephalosporins and aztreonam (dark blue), represented by cefotaxime (CTX), cefuroxime (CXM), ceftazidime (CAZ), aztreonam (AZT) and cefepime (CEP), higher generation of penicillins (light blue), represented by ampicillin (AMP) and piperacillin (PIP) and combinations of penicillins with beta-lactamases inhibitors (light violet), represented by ampicillin + sulbactam (AMS) and piperacillin + tazobactam (PIT); aminoglycosides (yellow) represented by gentamicin (GEN), amikacin (AMK), netilmicin (NET) and tobramycin (TOB); polymyxins (dark grey), represented by colistin (COL); combination of sulphonamides and trimethoprim (T/S, pink) and amphenicols (turquoise) represented by chloramphenicol (CMP).

### Characterizing the diversity of plasmid lineages across ST744

We detected plasmid replicons in the vast majority of ST744 genomes (872/892, 97.8%). The most prevalent large plasmid replicons included IncF (71.0% of strains), IncX (34.6%) and IncI1 (26.1%) (Fig. S3, Table S1), with 55.9% of strains carrying the FIB replicon, 30.7% the FIC replicon, and 29.1% the FII replicon, while the IncX1 plasmids were present in 20.4% of strains and IncX4 in 14.6%. The IncF plasmids, often containing multiple replicons, are known to have strongly contributed to the success and spread of multiple prominent ExPEC lineages^36,37^. Within ST744, we detected several notable and prevalent IncF plasmid lineages. This included F18:A:B1 plasmids (the most prevalent IncF plasmid replicon combination, detected in 129 genomes) (Fig. 2), which were distributed across both dominant serotypes and represented virulent ColV-like plasmids (Fig. S3, Table S1). We obtained a complete F18:A:B1 plasmid sequence from our reference isolate SGL242 (Fig. S4), which was 127131 bp length and carried the typical ColV plasmid-associated VAGs including *iroBCDEN*, *iucABCD*/*iutA*, *sitABCD*, *hlyF*, and *ompT*, without any ARGs. The second most dominant F plasmid type was F2:A:B (73 genomes), which was also detected across both clades and is not typically linked with ColV. ST744 genomes in which no plasmid replicons could be detected were mostly associated with *Livestock* in the UK (16/20 strains) (Fig. 2). In addition, we also observed a case where a Q1 plasmid was incorporated into the chromosome and the specific isolate (SGL280) carried no extrachromosomal plasmids. This genomic integration might have occurred for more strains, as the Q1 replicon was detected in 76.9% of strains (Fig. S3, Table S1).

### ST744 genomics and the link with ExPEC

To identify ST744-specific genomic regions and *ExPEC-*linked genomic regions in ST744 we used several complementary approaches. We first looked for the presence of insertion sequences (ISs) in ST744 (Table S10) and compared their prevalence to that of the general *E. coli* population (Table S10). ISs are commonly linked with genomic regions acquired via horizontal gene transfer, as they facilitate the transfer and further rearrangements of these regions^39,40^. For multiple ISs, we found that their prevalence is much higher in ST744 than across other *E. coli* genomes. Out of seven ISs identified in more than 90% ST744 genomes, four (IS*231*, IS*150*, IS*1202*, IS*903*) showed a large difference in prevalence (>50%) with the general *E. coli* population (Table S10). Upon checking the genomic regions linked to these ISs, we identified two potentially relevant regions, linked to IS*1202* and IS*150,* that contained several cold-shock genes that are not typically found in multiple dominant ExPEC STs (Table S11).

In our second approach, we leveraged our long-read-based assemblies of two reference genomes, SGL242 and SGL280. We compared these genomes with MG1655 to identify all regions of difference (RODs) between them (Table S12). The obtained ROD database was then used to evaluate the presence of the different RODs across both our ST744 collection and our *E. coli* species-wide genome collection (Table S13). This analysis identified a region (7727_AW) which was highly prevalent among ST744 (present in over 90% strains, Fig. 2, Table S12) and carried several genes that are strongly associated with *ExPEC* in general (threshold of 20% increase in prevalence in *ExPEC* genomes compared to *ND,* Table S13). In total, we identified 34 genes that displayed an increased *ExPEC* association, originating from 8 RODs (Table S13, Fig. 3B). These genes included *sitABCD*, encoding a metal (iron/manganese) transporter and a known VAG^41^, *fucP*, encoding a fucose permease, *mcmM*, a gene for microcin M production, several genes encoding patatin-like proteins (BHO22238.1, WWX45282.1), and a phosphotriesterase (XXW33867.1).

In a final, more holistic approach, we constructed an assembled pangenome reference for ST744, together with a gene-presence/absence table (Table S2). We filtered the pangenome reference (26,200 genes) to exclude genes that were present in the core genome of ST744 or only in a single strain, yielding a database of 16,226 genes. We subsequently screened this database against our *E. coli* species-wide genome collection to identify genes enriched in *ExPEC* genomes. This analysis resulted in the identification of 217 genes that displayed a high association with *ExPEC* (Table S14, Fig. 3B). Part of these genes were known VAGs (e.g., *fuyA* and *papB*), while others were linked to metabolism (e.g., *scrB*, *rhmD*), membrane transport (e.g. *alsC*, *fadL*), MGEs (e.g., IS*Ec53*, IS*200C*), and phage-related genes (WP_000514100.1, WP_049254721.1, WP_000683141.1, Table S14). Despite our extensive annotation efforts (automated annotation accompanied by manual curation of genes not receiving an automated annotation), a small portion of genes remained without any predicted function. However, some of these displayed a very high association with *ExPEC*, including one that differed by 42% in prevalence between *ExPEC* and *ND* genomes (Fig. 3B), suggesting they could play important roles in infection-related contexts (Table S14). Additionally, we checked which core genes of ST744 are not typical for *E. coli* overall (Table S15). From this analysis, lipopolysaccharide glucosyltransferase *waaG* and a putative fimbrial-like adhesin *ybdG* stood out as they were presented only in 5.6% and 9.6% of the EnteroBase (EB) collection, respectively. However, with involvement of *waaG* in LPS biosynthesis^42^, this probably reflects more substantial allele differences given our 90% threshold for identity rather than true gene absence.

To evaluate whether certain genes are associated with a specific group within the ST744 collection, we performed a genome wide-associations study (GWAS). When using a subset of the collection where the clinical relevance could be determined (500 strains), we observed that certain groups of genes are linked with ST744 strains specifically annotated as *ExPEC* (p < 0.0001, Table S16). These groups included virulence factors *(iucA*, *iutA*, *ompT*), colicins (*cma*, *cmi*, *cvaC*), large plasmids (*traY*, *traM*, *repB*), other mobile genetic elements (*ISEsa1*, *yhgA*), toxin-antitoxin systems (*pemI*, *vapB*) and antimicrobial resistance genes (*bla*_CTX-M_, *ant1*). Many of these genes represent a backbone of, or are typically carried by, large F plasmids. Accordingly, we also observed that strains of H9 serotype tend to carry F plasmids more frequently and they have higher representation of *ExPEC* strains (Fig. 3A).

### Phenotypic characterization of *E. coli* ST744 identifies growth differences in specific environments

As our genomic analysis identified specific ST744 characteristics that might lead to differences in certain phenotypes, we set out to test some of these directly. For this, we selected a subset of 12 ST744 strains from our collection, which we combined with three representative strains of other ExPEC STs (ST69, ST155, and ST354) and the MG1655 laboratory strain for comparison. To assess whether the presence of additional cold-shock proteins in the ST744 strains, which were absent in ST155 and ST354, would enhance their growth at lower temperatures, we first measured the growth of these strains at 37°C and 25°C (Fig. 5AB, Fig. S5A). While growth rates that were extracted from the obtained growth curves revealed no significant differences at 37°C, all 12 ST744 strains and the three representative strains from other STs displayed elevated growth rates at 25°C compared to the MG1655 lab strain (Fig. 5AB, Fig. S5A). This finding suggests a potential growth advantage of ExPEC strains, or at least the included subset, at lower temperatures, yet also indicates that an increased presence of cold-shock genes does not underlie this phenotype (as these genes are absent in ST155 and ST354 but present in MG1655; Table S17).

**Figure 5.**
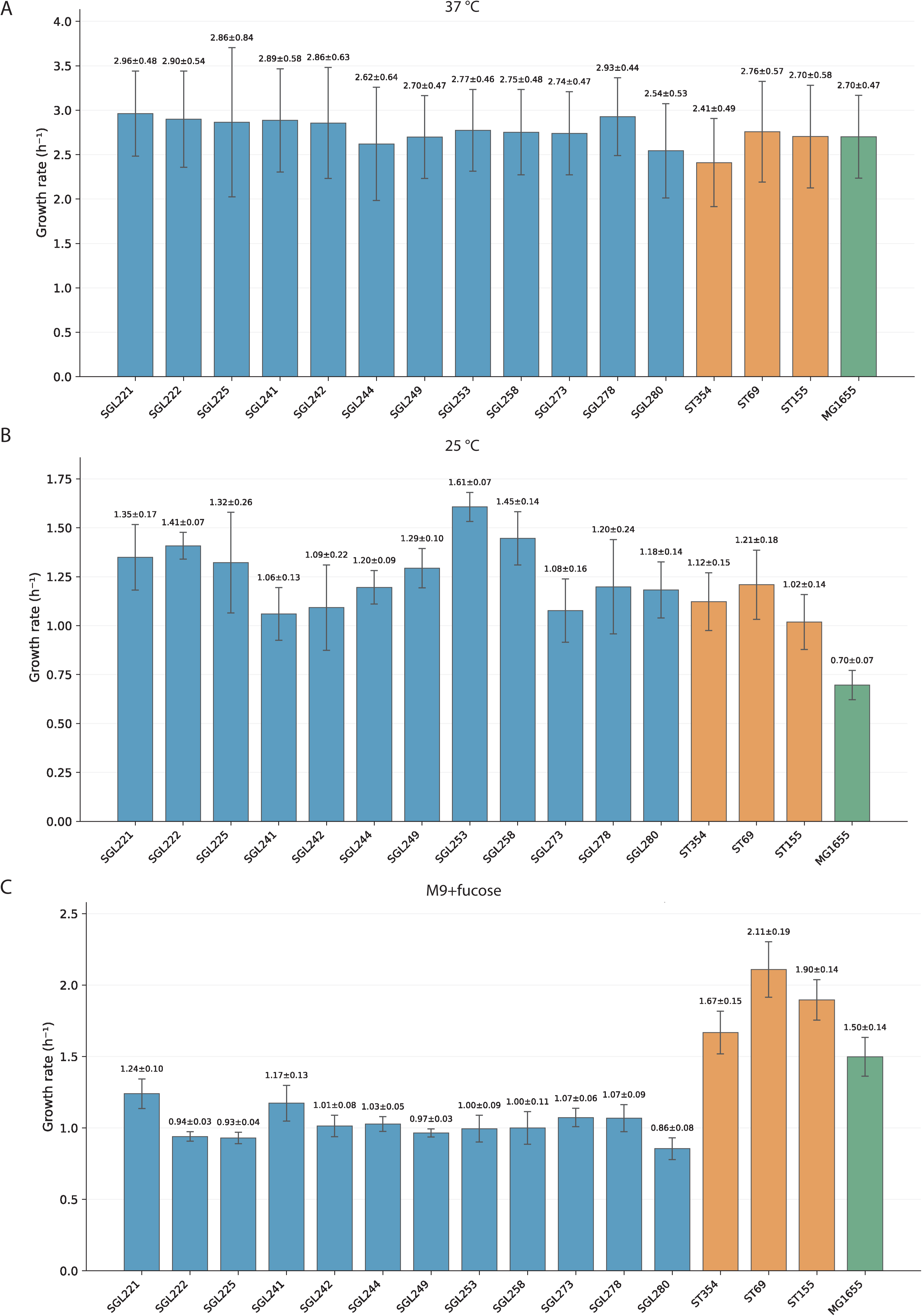
Growth rates comparison. Comparison of growth rates (h-1) for 12 ST744 strains (blue), 3 comparative strain of other STs (orange) and MG1655 (green) for growth on LB at 37°C (A), at 25°C (B) and on M9 with 0.2% L-fucose (C). Error bars indicate mean +-standard deviation.

To determine a potential role for the fucose permease, FucP, we also grew the same set of strains on minimal defined medium containing 0.2% of L-fucose as a carbon source (Fig. S5B). However, across our tested isolates, we observed that the fucose-supported growth rate was lower for ST744 than for the other STs or MG1655, suggesting *fucP* contribution to fucose utilization is likely limited (at least in these laboratory settings) (Fig. 5C), especially given that two of the tested ST744 strains lacked the *fucP* (SGL241 and SGL253, Table S17).

### Image-based profiling of *E. coli* ST744 reveals extensive within-clade diversity and establishes genotype-to-phenotype connections at the cellular level

To provide additional insight into the within-clade variability of ST744, we performed a microscopic characterization of 32 ST744 strains (while also including a MG1655 laboratory strain as control). For this, we grew the strains in LB and stained exponentially growing cells with the DNA dye DAPI to visualize their nucleoids. We subsequently imagedthe cells usingphase contrast and fluorescence microscopy (Fig. 6A), and used our BactoScoop pipeline^43^ to extract single-cell level features related to cell and nucleoid morphology. Already visually, we observed clear differences in the appearance of specific strains (Fig. 5A). For example, SGL221 cells were wider, SGL226 cells longer, SGL236 cells shorter, and SGL254 cells larger (longer and wider) (Fig. 6A). In addition, ST744 strains differed in their nucleoid dynamics, with nucleoids segregating earlier/later in the cell division cycle of specific strains, as revealed by demographs of their DAPI signal (Fig. 6B). We further confirmed this by calculating the relative timings of nucleoid separation, nucleoid constriction, and cell constriction of these strains using the relative timing inference method^44–46^ (Fig. 6C).

**Figure 6.**
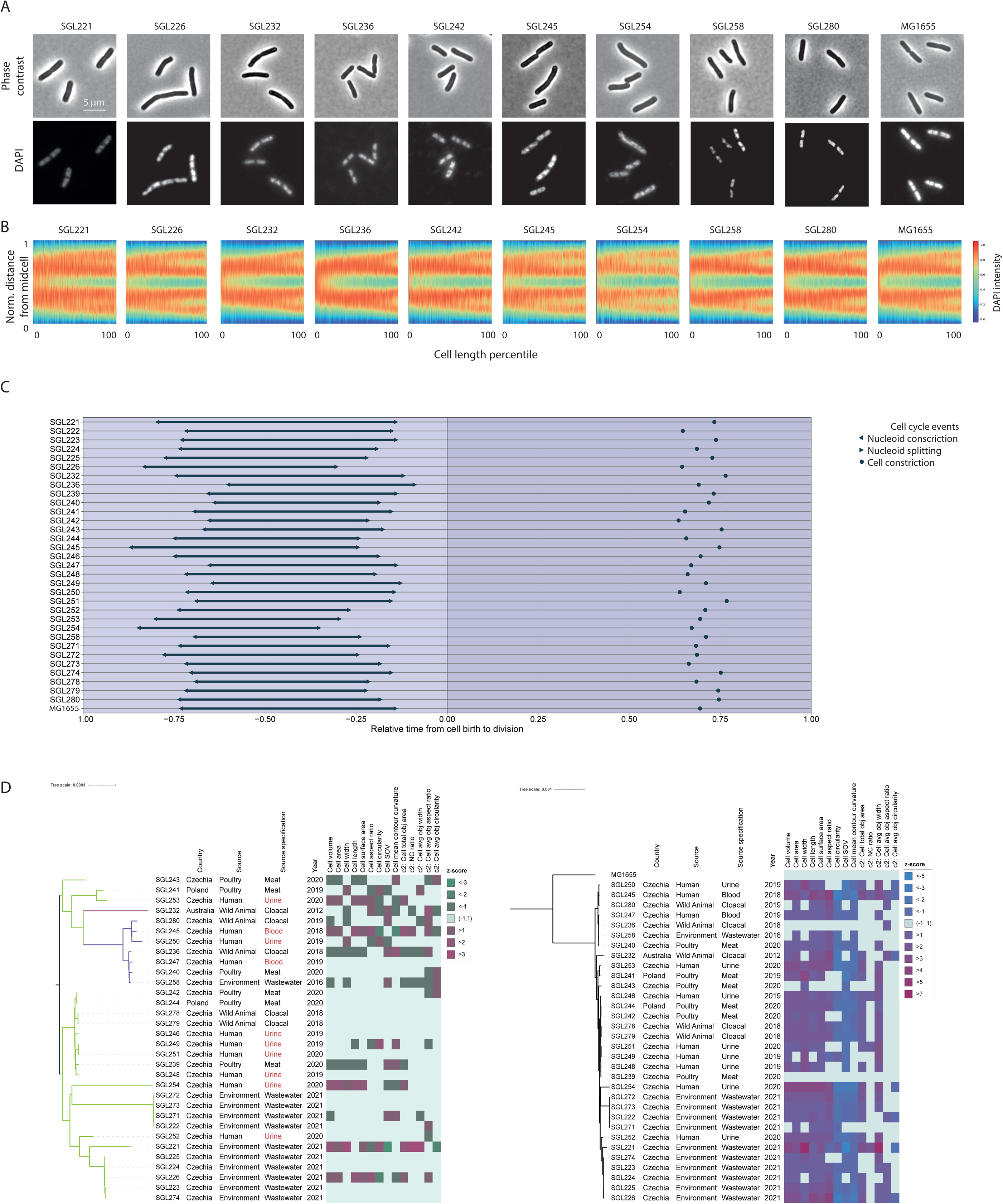
Image-based profiling of *E. coli* ST744. A) Microscopy images of selected ST744 and MG1655 for the phase channel and DAPI channel. Strains identifiers are indicated above the phase figures. B) Demographs of DAPI signal intensity showing the nucleoid dynamics during exponential growth. DAPI signal intensity is plotted for each cell (sorted by cell length) along the normalized cell length, with 0 and 1 indicating cell poles. C) Cell cycle timing visualization across two cell division cycles, as some events happen in the previous or next cell cycle. D) Phylogenetic tree for 32 ST744 strains alone (left panel) and compared to MG1655 (right panel), accompanied by 14 representative phenotypic features extracted from the microscopy data and visualized as a z-score.

Using our BactoScoop image-analysis pipeline, we extracted >300 quantitative features from our microscopy images (Table S18). From this, we selected 14 biologically interpretable features that captured broad variations in cell and nucleoid morphology. For each of these features, we calculated z-scores for each strain based on the mean value for all strains (ST744 isolates + MG1655; Fig. 6D, Table S19) and based on the mean value of the ST744 isolates (to better emphasize differences between them) (Fig. 6D, Table S20). This analysis captured initial qualitative observations (e.g., wide SGL221 cells) and revealed extensive diversity, both in comparison to MG1655 and among ST744 strains, for almost all included phenotypic features (Fig. 6D). Remarkably, SGL221, which displayed the most pronounced deviations from other ST744 strains, also displayed considerable phylogenetic differences (with 236 unique SNPs being detected within the core genes and 71 accessory genes being linked uniquely to this strain).

Large phenotypic differences were not always associated with large genomic differences. For example, strains SGL223-226 are clonal, yet a single genomic region consisting of eleven genes (i.e., *yedP*, *yedQ*, *yodC*, *yodD*, *yedA*, *vsr*, *dcm*, *yedJ*, *drpB*, *dsrB* and *dgcQ*) was missing in SGL226 (with no unique SNPs being linked to this strain specifically). While most of these genes are so-called y-genes of which the exact function of is not known, DprB has been identified as a nonessential cell division protein in *E. coli*^47^ and its deletion could thus underlie the elongated cell phenotype seen in this strain (and not in SGL223-225; Fig. 6D). Similar phenotypic differences between closely-related strains were observed for a pair of urinary pathogenic *E. coli* (UPEC) isolates (SGL249 and 251, 4 SNPs), meat isolates from *Poultry* (SGL242 and SGL244, 0 SNPs) and a clade of *Wastewater* isolates (SGL222, SGL271, SGL272 and SGL273) (Fig. 6D). The UPEC clonal pair differed in the presence of over 200 accessory genes (202 additional genes for SGL251 and 40 for SGL249) (Table S2). The pair from *Poultry* differed in presence of 36 genes (18 each) and 52 accessory genes were detected for the *Wastewater* clade which were not present in all four but were detected in at least one strain from the *Wastewater* clade (Table S2).

## Discussion

Our species-wide analysis places the genomic properties of individual ExPEC lineages within the broader diversity of *E. coli* and reveals a negative relationship between the average numbers of recognized VAGs and ARGs across the 70 most common ExPEC STs. Within this landscape, ST744 exhibited the highest average ARG content, motivating a detailed characterization of its population structure, genomic composition and phenotypic properties. By combining comparative genomics with antimicrobial susceptibility testing, growth measurements and image-based profiling, we show that a broadly-shared multidrug-resistant background coexists with considerable genomic and cellular diversity within this lineage. Together, these findings provide an in-depth characterization of *E. coli* ST744 while also illustrating both the value and the limitations of using genomic information to infer phenotypic properties.

Our inverse relationship between VAG and ARG content extends previous observations from individual lineages and smaller strain collections^17,26,27,48,49^ to a broader comparison of common ExPEC STs. This pattern is consistent with the contrasting genomic profiles of the three main ExPEC groups (i.e., *Classical high-virulence ExPEC*, *Emerging MDR ExPEC*, and *One-Health high-risk ExPEC*). While our observation likely reflects a major evolutionary transition in the ExPEC population, given that each main group emerged at a different point in time, it is important to emphasize that it does not establish causality and therefore does not provide evidence for a direct trade-off between resistance and virulence. Moreover, the number of recognized VAGs is an imperfect proxy for pathogenic potential, as individual determinants differ in their contribution to infection and their effects can depend on expression, genetic background and host context, and specific additional virulence traits might not have been discovered yet. This is especially relevant for *One-Health high-risk ExPEC*, which often carry comparatively few recognized VAGs despite increasingly being involved in urinary tract, bloodstream and other extraintestinal infections^21,50,51^. An example of this are the large differences in virulence that have been observed between two clades of ST410, the best studied *One-Health* ExPEC ST, that have similar VAG counts^21^. In a *Galleria mellonella* infection model, a clade of B5/H24RxC (median of 176 putative VAGs) led to a significantly lower survival rate (and thus higher virulence) than a comparative clade of B4/H24RxC (median of 166 VAGs), even surpassing that of a hypervirulent *Klebsiella pneumoniae* strain K1088^21^. Likewise, ARG counts do not always capture the breadth or magnitude of phenotypic antibiotic resistance^52^. While genomic properties should thus be interpreted with care, ST744 remains a remarkable lineage given its extreme levels or ARGs.

ST744 is a member of PG A and clonal complex 10, and is closely related to ST10, which is considered a notable opportunistic ExPEC ST^4,53^. The first ST744 strain recorded in EB came from a human sample in France in 2006. One of the earliest mentions of ST744 in the literature came from sampling in 2008-2009, after which a single ESBL-producing isolate of ST744 was isolated from a human in Laos^54^. ST744 is commonly reported in studies focused on important antibiotic resistance determinants, including mainly plasmid-mediated colistin resistance (*mcr* genes), ESBL and AmpC, and resistance to fluoroquinolones. ST744 isolates carrying these determinants were reported from the environment and surface water and wastewaters^55–57^, wild birds^58–63^, other wild animals^64,65^, poultry^66–73^, other livestock^65,74–79^, companion animals^80,81^, and humans^29,54,66,82^. Previous studies have also documented the pathogenic potential of ST744, despite carrying a comparatively limited repertoire of recognized VAGs relative to *Classical ExPEC* and *Emerging MDR* lineages. In a global analysis of >10,000 ExPEC blood stream infections (BSIs), ST744 was identified as a prevalent causative agent associated with MDR^83^, and patients with BSIs caused by ST744 were reported from Denmark^66^, South Africa^84^, the Czech Republic^29^ and China^85^. In addition, ST744 has been implicated in UTIs in different countries^28,29,80,86^. Infections caused by ST744 are not limited to humans but occur in animals as well, including in a dog with a prostatic abscess^87^, dogs with UTIs^88^, and diseased livestock^89,90^. The identification of closely related isolates from different sources, sometimes collected in the same country and year, is characteristic of a *One-Health* lineage and consistent with circulation across connected human, animal, and environmental settings, or exposure to shared sources. Moreover, ST744 is dispersed across the globe, given it has been detected in >50 countries across all continents (with the exception of Antarctica)^31,32,63,84,91,92^. Building on all these observations, our analysis resolved the global population structure of an expanded collection of 892 ST744 genomes and placed this diversity in a species-wide context.

A notable feature of almost all ST744 strains is the distribution of resistance determinants across chromosomal mutations, integrated resistance regions and variable accessory elements. Nearly all genomes in our collection carried the common fluoroquinolone resistance-associated point mutations (99.1%), and detection of ESBL/AmpC genes and *mcr* genes was relatively high (49.6% and 12.7%, respectively). Moreover, we detected a carbapenemase gene in 7.0% of genomes, with *bla*_NDM-1_, *bla*_OXA-48_ and *bla*_NDM-5_ being the most commonly found. In this regard, ST744 follows a similar trend as ST167 and ST410, which both have been previously associated with multiple carbapenemases as well^10,50,51^. While many *E. coli* lineages depend on plasmids for obtaining antibiotic resistance, a large proportion of ST744 contains a chromosomal region carrying *intI1* integrase, mercury resistance markers, and a variable amount of ARGs (5 and 10 ARGs in the case of our two long-read sequenced strains). The physical linkage between antibiotic and mercury resistance genes raises the possibility of co-selection in environments containing elevated metal concentrations, where the presence of mercury in the environment favors maintenance of the linked ARGs (even in the absence of direct antibiotic selection), similar to what has been proposed for other antibiotic resistance-associated genetic elements^93–95^.

The two dominant ST744 clades, associated with O101:H9 and O101:H10 serotypes, provide a framework for examination of genomic variation within the lineage. Against a broadly shared resistance background, variation in accessory gene content, plasmid carriage and clinical annotation suggests that ST744 contains subpopulations with potentially different biological properties. The stronger association of *ExPEC* annotations with O101:H9 isolates and their increased IncF plasmid carriage (Fig. 3A) raises the possibility that accessory virulence-associated functions contribute to these differences. This is further supported by the complete sequence of the F18:A:B1 plasmid, the most common F plasmid replicon combination in our collection, revealing important VAGs but no ARGs. Together, these findings suggest that F plasmids may provide functions relevant to pathogenic potential within a lineage with a relatively limited repertoire of recognized chromosomal VAGs but that is already antibiotic-resistant. In contrast, F plasmids have frequently contributed to the acquisition and dissemination of AMR determinants in successful *Emerging MDR ExPEC* lineages, like ST131^96^.

Our analyses also identified >200 genes in the ST744 accessory pangenome that were enriched among *ExPEC* genomes across the species-wide collection. These genes included obvious candidates such as known VAGs, but also metabolism-and membrane transport-related genes. This broad functional distribution suggests that conventional VAG catalogues may capture only part of the genetic repertoire associated with extraintestinal infection. Metabolic and transport functions could support colonization, competition or persistence, thereby contributing to infection without acting as dedicated virulence factors. Previous work has similarly implicated metabolic versatility in the success of ExPEC lineages and in the regulation of virulence-associated factors^53,97^. However, and this is true for all associations in our study, enrichment does not demonstrate a causal contribution to infection. Associations may arise through shared ancestry or linkage to other determinants, and genomes of undetermined clinical status are not a verified non-pathogenic control group. Future functional studies are always required to determine the phenotypic consequences of harboring specific genes.

Our phenotypic characterizations highlight that the interpretability of genomic information depends on the trait being considered. For antibiotic resistance, established genetic determinants generally corresponded with susceptibility measurements. This included agreement between fluoroquinolone resistance and the relevant chromosomal mutations, between colistin resistance and the presence of *mcr-1*, and between resistance to extended-spectrum beta-lactams and the presence of an ESBL/AmpC. This correspondence was not universal however, as illustrated by several isolates carrying trimethoprim/sulphamethoxazole resistance genes without displaying the expected resistant phenotype. Nevertheless, the overall agreement between genotype and phenotype supports the value of established resistance determinants for predicting susceptibility to the antibiotics tested.

In contrast, the growth consequences of candidate metabolic and stress-associated genes were less readily inferred from their presence. While ST744 isolates grew faster than MG1655 at 25°C, this pattern could not be explained by the presence of additional cold-shock genes, as representative strains of other ExPEC STs lacking these genes also grew faster. Likewise, the presence of the fucose-associated region did not translate into enhanced growth on fucose under the conditions tested. An explanation for these apparent discrepancies between genotype and phenotype likely lies in the fact that growth represents a complex phenotypic trait that results from the combined activity and regulation of many cellular processes. As such, the presence of specific metabolic and/or stress-associated genes likely exerts a less straightforward phenotypic effect than antibiotic resistance genes or mutations. In addition, our laboratory setup differs significantly from realistic infection environments, complicating a physiologically relevant comparison. Moreover, fucose is also known to contribute to infection by functioning as a signalling molecule promoting biofilm formation^98^, influencing the expression of virulence factors^99^, and affecting colonization efficiency^100^. These results emphasize the importance of experimentally distinguishing, in realistic settings if possible, plausible functional annotations from demonstrated phenotypic consequences.

Quantitative microscopy further revealed extensive variation in cell morphology, nucleoid morphology, and cell cycle timings within ST744, including among clonal isolates. These findings demonstrate that close genomic relatedness does not necessarily imply equivalent cellular phenotypes. The elongated phenotype of SGL226, which lacks an eleven-gene region present in closely related isolates, provides a particularly informative example. This deletion includes *drpB*, encoding a nonessential cell division protein^47^, making it a plausible contributor to the phenotype. However, the involvement of other deleted genes or their combined effects cannot be excluded. More generally, phenotypically divergent near-clonal isolates provide opportunities to prioritize candidate determinants that underlie the observed divergence.

Although phenotypic characterization is widely employed in ExPEC research, most studies focus on traits directly associated with antimicrobial resistance, pathogenicity or adaptation to environmental stress (e.g., growth under antibiotic exposure, biofilm formation, adhesion, invasion and resistance to host-associated stressors)^101,102^. In contrast, fundamental cellular properties, including cell morphology and nucleoid organization, have received comparatively little attention. Nevertheless, these features reflect essential aspects of bacterial physiology that could contribute to colonization, persistence or pathogenicity of specific strains. While this contribution cannot be inferred from our experiments directly, integrating quantitative cellular phenotyping with comparative genomics provides a means to identify and investigate physiological diversity within clinically relevant lineages. Testing candidate genetic differences and determining whether the observed phenotypes persist across environments will help establish their mechanistic basis and biological significance. Future studies should therefore investigate the stability of the observed phenotypic differences across environmental conditions, identify candidate genetic determinants through detailed comparative genomic analyses, and evaluate whether specific cellular characteristics are associated with traits relevant to host colonization, persistence or pathogenicity. Ultimately, a better understanding of the relationship between fundamental bacterial cellular phenotypes and pathogenic behavior could provide additional insights into the biological diversity and adaptive potential of ST744 and other ExPEC lineages.

Overall, our study places ST744 within the species-wide genomic diversity of *E. coli*, identifying it as a multidrug-resistant lineage that occurs across human, animal, and environmental sources. Beneath its broadly shared MDR profile, ST744 displays considerable variation in accessory gene content, resistance-region organization and plasmid carriage. This genomic diversity is accompanied by extensive variation in cellular properties, even among closely related isolates, highlighting the biological heterogeneity concealed within a single sequence type. The differing correspondence between genomic information and antibiotic resistance, growth and (sub)cellular phenotypes further underscores the value of combining comparative genomics with experimental characterization to provide more complete understanding of ST744 and a framework for investigating how genetic variation translates into biological diversity within other ExPEC lineages.

## Methods

### Strains and ST744 genomic data collection

For phenotypic characterization, a total of 32 *E. coli* ST744 isolates were included in this study. These strains originated from different sources and countries (albeit mostly from within Czechia) and were obtained via our previous studies^29,31,55,63,79,103^. The metadata for all of them are specified in Table S1. Lab strain MG1655 was used as a comparative control strain for several experiments and analysis.

Three strains of different ExPEC STs (ST354, ST69 and ST155) were used for comparison in growth experiments as they differed in the presence of the targeted genes.

For the genomic data collection, we downloaded all ST744 genomes (Table S21) which were accompanied by source-related metadata in June 2023 from EB (https://enterobase.warwick.ac.uk/species/index/ecoli). We filtered out genomes with insufficient coverage (<30x) to obtain a final collection of 860 genomes from EB, which was combined with the genomes of the 32 ST744 strains from our lab collection to yield the final genomic collection of 892 strains (Table S1). We provided additional annotation to the genomes and labelled them as *ExPEC* in case they came from an ExPEC-relevant source (e.g., urine, blood) or *Clinical* in case the source had clinical relevance (e.g. patient, hospital).

### Repurposing of large-scale genomic collection for specific analyses

We previously compiled a large-scale genomic collection of 131,610 *E. coli* genomes, from which 20523 were *ExPEC*-relevant^7^. The extent of their virulence-associated genes and antibiotic resistance genes was already evaluated in our previous study using VFDB (http://www.mgc.ac.cn/VFs/)^104^ and ResFinder (v.3.0)^105^ databases. We used the outputs from VFDB and ResFinder to calculate the average VAGs and ARGs, respectively, for the 70 dominant ExPEC STs.

The 70 dominant ExPEC STs represented 82% of all *ExPEC* genomes and were represented by a minimum of 35 *ExPEC* genomes. The collection was also used to determine which of the ST744-associated genes might be more broadly associated with more specific *E. coli* categories (e.g., *ExPEC*-associated). Such screens were performed using ABRicate (v1.4.0, https://github.com/tseemann/ABRICATE) with 90% thresholds for query coverage and identity.

### Phylogenetic analysis

The phylogenetic trees produced within this study include a tree for the complete ST744 collection (892 genomes), our strain collection (32 genomes) and our strain collection compared to MG1655 (33 genomes). The phylogenetic analysis was based on Prokka (v1.14.5) open reading frames annotation^106^, and multi-fasta alignment with Roary (v3.7.0)^107^. RAxML^108^ with GTRGAMMA, supported by 1,000 bootstraps, was used to calculate the trees. SNP distance across all pairs of genomes (Table S3) was evaluated using snp-dists (v. 0.7.0; https://github.com/tseemann/snp-dists) with 10 SNPs being considered as a threshold for clonality^109^ We examined the set for clonal pairs from the same *Country* and *Year*, yet coming from different *Niches* to emphasize the One-Health potential of ST744 and highlight the possible epidemiological link between strains from different sources (Table S4).

### Comparative genomics and GWAS

Multiple approaches were used to explore the genomic content of ST744, in comparison with the general *E. coli* population, or to explore specifics of different groups (e.g., *ExPEC* genomes within ST744). The preliminary exploratory approach used screening of ISs using ISfinder^110^within our ST744 collection (Table S10) and compared it with a previous dataset of general population ISs distribution using a collection of 1,933 complete *E. coli* genomes (downloaded on NCBI on 01/02/2022 as all available complete genomes, Table S22). We further focused on ISs which were highly prevalent (>90%) in ST744 and at the same time, considerably less prevalent (min. 50% difference) in the general *E. coli* population. We explored annotated contigs containing these ISs in ST744 using Geneious (v7.1.9) and identified that IS*1202* and IS*150* are part of two genomic regions linked with additional cold-shock proteins, which are not typically present in other typical ExPEC STs (Table S11).

In our second approach, we evaluated RODs between our two long-read sequenced strains (SGL242 and SGL280) and MG1655 using alignment with progressiveMauve^111^. A total of 30 and 12 regions containing 520 and 184 predicted genes were obtained for SGL242 and SGL280, respectively (Table S12). These genes were used to create a customised database, that was subsequently used to run in ABRicate with thresholds of 90% query coverage and identity against our large genomic collection of 131,610 *E. coli*. From these results, we determined genes that are enriched in *ExPEC* (at least 20% difference between *ExPEC* and *ND* category), which are summarized in Table S13. From these, we focused for subsequent growth experiments on *fucP*, which had over 23% higher prevalence in *ExPEC*.

As the phylogenetic analysis provided a table with all predicted genes within our set, our broadest approach to estimate which accessory genes of ST744 are linked with general *E. coli* and ExPEC population was based on this dataset. The database included a total of 16,226 predicted genes, which were present in maximally 891 genomes (core genes were not included) and minimally 2 genomes. The screen was based on ABRicate, using the same thresholds of 90%. With this approach, we identified a total of 217 genes that are enriched in *ExPEC* (with a threshold of 20% to the comparative group). We also screened the *E. coli* genomes for the core genes of ST744 to determine which of these are atypical (<10%) in the general *E. coli* population (Table S15).

To explore genomic links within the ST744 and specific groups including *Serotypes*, *Niche* or *ExPEC* status, we performed genome-wide association study using gene_presence_absence file generated by Roary^107^ as an input for Scoary^112^. For the ExPEC evaluation, we used a limited dataset of 500 genomes where the clinical status could be assigned based on additional source information.

### Screen using established databases

We always used ABRicate with a 90% threshold for both query coverage and identity to screen for the presence of VAGs using VFDB^104^, ARGs using ResFinder (v.4.4.1)^105^, plasmid replicons using PlasmidFinder^113^ (v3.0.3). AbriTAMR (v1.0.18)^114^ was used to determine point mutations relevant in antibiotic resistance. pMLST was used to determine replicon sequence type (RST) of F plasmids and pMLST of other plasmids^113^. AbriTAMR^114^ was used to screen for heavy metal resistance.

### Phenotypic assessment of antimicrobial resistance

The antimicrobial susceptibility of our 32 isolates was tested for 24 different antimicrobial compounds or their specific combinations by broth microdilution method using the MIKRO-LA-TEST MIC GN 1, MIC GN 2 kit suitable for Gram-negatives according to the manufacturer’s instructions (Erba Lachema, Brno, Czech Republic). The EUCAST (2024)^115^ clinical standards were applied to determine whether an isolate is resistant, susceptible, or showing an intermediate phenotype for 20 out of the 24 compounds as the rest do not have any EUCAST breakpoints anymore.

### Growth experiments evaluating the impact of cold stress and a specific carbon source

For growth experiments, a total of 12 ST744 strains were selected together with 3 comparative strains of other STs (ST69, ST155, ST354) and an MG1655 control strain. All strains were inoculated in 200 µl LB broth in a 96-well plate and incubated overnight (16-20h) with shaking (200 RPM) at 37°C. The following day, cultures were diluted by 1:10,000 in 200 µl LB (temperature test) or M9 minimal medium with 0.2% L-fucose and incubated at 37°C or 25°C (for the cold stress test), shaking at 200 RPM in a Synergy HT Multi-Mode Microplate Reader (Biotek, USA) with measuring OD600 every 15 minutes for 24 hours. All experiments were done in biological and technical triplicates and negative controls (wells with medium but no inoculation) were included in each experiment. Growth curves were generated in R using ggplot2 from blank-corrected (well-specific blank) OD600 measurements, with three technical replicates averaged within each of three independent biological replicates. The curves represent the mean OD600 over time, with shaded areas indicating ±1 standard deviation across biological replicates (Fig. S5). Growth rates were determined from smoothed, log₂-transformed OD_600nm_ measurements. Multiple candidate exponential growth phases were identified by fitting linear regressions to sliding windows of 7 consecutive time points across the entire growth curves. The regression corresponding to the actual exponential phase was selected based on three criteria: a minimum increase in OD₆₀₀_nm_ of 0.002 between the starting and ending time points, a high goodness of fit (R² > 0.98), and the highest slope. To avoid constraining the analysis to the initially defined window size, the selected exponential phase was subsequently extended towards both earlier and later time points whenever the inclusion of additional data points remained consistent with exponential growth (within the limit of increasing the average residuals by 3, starting from the first fit). This iterative extension was intended to improve the robustness of the final growth rate estimate, corresponding to the slope of this regression in h⁻¹.

### Fluorescence microscopy

All strains were imaged by fluorescent microscopy in biological triplicates. Every sample was diluted 1:10,000 from an overnight preculture (3 ml LB, 37°C, 200 RPM shaking) and grown in 3 ml LB in sterile glass tubes into exponential phase (OD600 approx. 0.1). When the required OD was reached, 500 μL was mixed with 0.5 μL of DAPI (4′,6-diamidino-2-phenylindole, Sigma-Aldrich, Germany) and incubated for another 15 minutes at 37°C, 200 RPM to stain the DNA. Subsequently, 0.5 μL of cells with DAPI were spotted on 1.0% UltraPure™ Agrose (Invitrogen, Thermo Fisher Scientific, UK) pads supplemented with M9 medium and covered with a cover glass. Microscopy was performed using a Nikon Eclipse Ti2-E inverted epi-fluorescence microscope, Plan Apo λ 100× DM oil objective (NA = 1.45; Nikon, the Netherlands), Ph3 condenser annulus (Nikon, the Netherlands), and a Kinetix A22J723021 sCMOS camera (Teledyne Photometrics, USA). NIS-Elements AR software was used to control and set up the microscopy settings. We used a pE-800 light source (CoolLED, UK) with eight separate LED channels allowed narrowband excitation, and filtercubes with BrightLine Pinkel filtersets (penta-band set: LED-DA/FI/TR/Cy5/Cy7-5X-A; Semrock, IDEX Health and Science, USA) and OptoSpin filterwheels (Cairn GmbH, Germany) for fluorescence excitation with following conditions: 400 nm excitation, excitation filter:FF01-378/52, dichroic FF409/493/573/652/759-Di01, emitter filter cube: FF01-432/515/595/681/809, emitter filter wheel: FF01-432/36). For every sample, we aimed to capture 20 non-overlapping positions to obtain a sufficient number of single cells.

### Image analysis and feature processing

Phase-contrast images were segmented using Omnipose^116^, followed by mesh construction, SVM-based curation^117^ and feature extraction using BactoScoop (v.0.1.2)^43^. Cell morphology, DNA fluorescence profiles and nucleoid features were extracted from curated cells using the ‘morphological’, ‘profiling’, and ‘objects’ modules. For z-score analysis, features were summarized as medians within each strain-replicate combination. Replicate profiles were standardized using the mean and standard deviation of wild-type MG1655 (SGL315) replicate profiles, and the resulting z-scores were averaged per strain. Separately, replicate medians were averaged per strain and standardized across the non-wild-type strain population.

### Cell cycle timings

Cell cycle timings were estimated using an established calculation method based on the fractions of the total cell population in the different cell cycle stages and assuming binary fission^44,46,118^. Cells with a maximal indentation in the DNA signal profile (feature c2_object_avg_consDegree) exceeding 0.2 A.U. were considered to be undergoing detectable nucleoid segregation, whereas cells with a mid-cell decrease in cell width (feature cell_constriction_degree_width_um) exceeding 0.1 µm were considered to be undergoing detectable cell division. These threshold values were conservatively selected by examining the degree of constriction of individual cells from each replicate, plotted as a function of cell length which was used as a proxy for cell age. The numbers of nucleoids at birth and at division were estimated based on the distribution of nucleoid numbers per cell in the population. This estimation relied on the assumption that most cells would have a nucleoid number corresponding to that observed at birth, given that an exponentially growing cell population is expected to contain twice as many newborn cells as cells approaching division. These values were then used to assess the degree of overlap between successive DNA replication cycles.

### Statistical analysis and data visualization

We used RStudio (v2023.06.0) with R (v4.4.1) and the packages ggplot2, dplyr, tidyr, corrplot, viridis, grid, ggplotify, RColorBrewer, and polycor to merge ABRicate results from all our and established databases with the metadata, summarize the results, perform statistical analysis and create individual plots. Density patterns for VAGs and ARGs distribution (Fig. S1) were based on a kernel density estimation using VAGs and ARGs numbers for our whole EB collection (131610 genomes, Fig. S1A), *IPEC*-labelled genomes (Fig. S1B) and *ExPEC*-labelled genomes (Fig. S1C). Spearman correlation was used to estimate the relation between VAGs and ARGs content of dominant ExPEC STs. Python (v3.10) was used for microscopy image analysis, growth data analysis and cells cycle timing analysis. The code is specified and explained at https://github.com/Govers-Lab/Nesporova_et_al_2026.

We used Geneious (v7.1.9) to check annotations and alignments of specific genomic regions. The complete sequence of F18:A:B1 plasmid from SGL242 was visualized using HERA^119^. The phylogenetic trees were visualized using the Interactive Tree of Life (iTOL v6)^120^.

## Declarations

### Ethics approval and consent to participate

Not applicable.

### Consent for publication

Not applicable

### Availability of data and materials

All data generated or analyzed during this study are included in this published article and in supplementary information files. The genomic data are available via BioSample ID (NCBI) or UberStain ID (EnteroBase) specified in Table S1. The code is available at https://github.com/Govers-Lab/Nesporova_et_al_2026.

### Competing interests

The authors declare that they have no competing interests

### Funding

This work was funded by the European Union under Horizon Europe: MSCA Actions (Project 101105027) and by junior postdoctoral fellowship from the Research Foundation – Flanders (file number 1251624N) awarded to K.N. This work was further supported by a start-up grant (STG/21/068) from the KU Leuven Research Fund awarded to S.K.G and by Czech Science Foundation (24-12527S) to M.D.

### Author contributions

Conceptualization, K.N., M.D. and S.K.G.; methodology, K.N., M.D. and S.K.G.; software, K.N., B.S. and A.M.; formal analysis, K.N., B.S., A.M., A.V., J.P., K.K.; investigation, K.N.; data curation, K.N., B.S.; writing – original draft, K.N. and S.K.G.; writing – review & editing, K.N., M.D., B.S., A.M., A.V., J.P., K.K.; and S.K.G.; visualization, K.N., B.S and A.M.; supervision, S.K.G. and M.D.; project administration, S.K.G.; funding acquisition, K.N., M.D. and S.K.G.

## Acknowledgements

We would like to thank the members of Govers laboratory for valuable feedback on the manuscript. We acknowledge all EnteroBase contributors. The computational resources and services used in this work were provided by the VSC (Flemish Supercomputer Center), funded by the Research Foundation Flanders (FWO) and the Flemish Government department EWI. K.N. was funded by a junior postdoctoral fellowship from the Research Foundation—Flanders (file number 1251624N). This work was also supported by a start-up grant (STG/21/068) from the KU Leuven Research Fund and by the European Union (ERC Starting Grant BacterialBlueprint 101040800).

## Notes

### Competing Interest Statement

The authors have declared no competing interest.

