## Supplementary material for "From species-wide genomics to cellular diversity: characterizing multidrug-resistant extraintestinal pathogenic *Escherichia coli* ST744": Figure S1

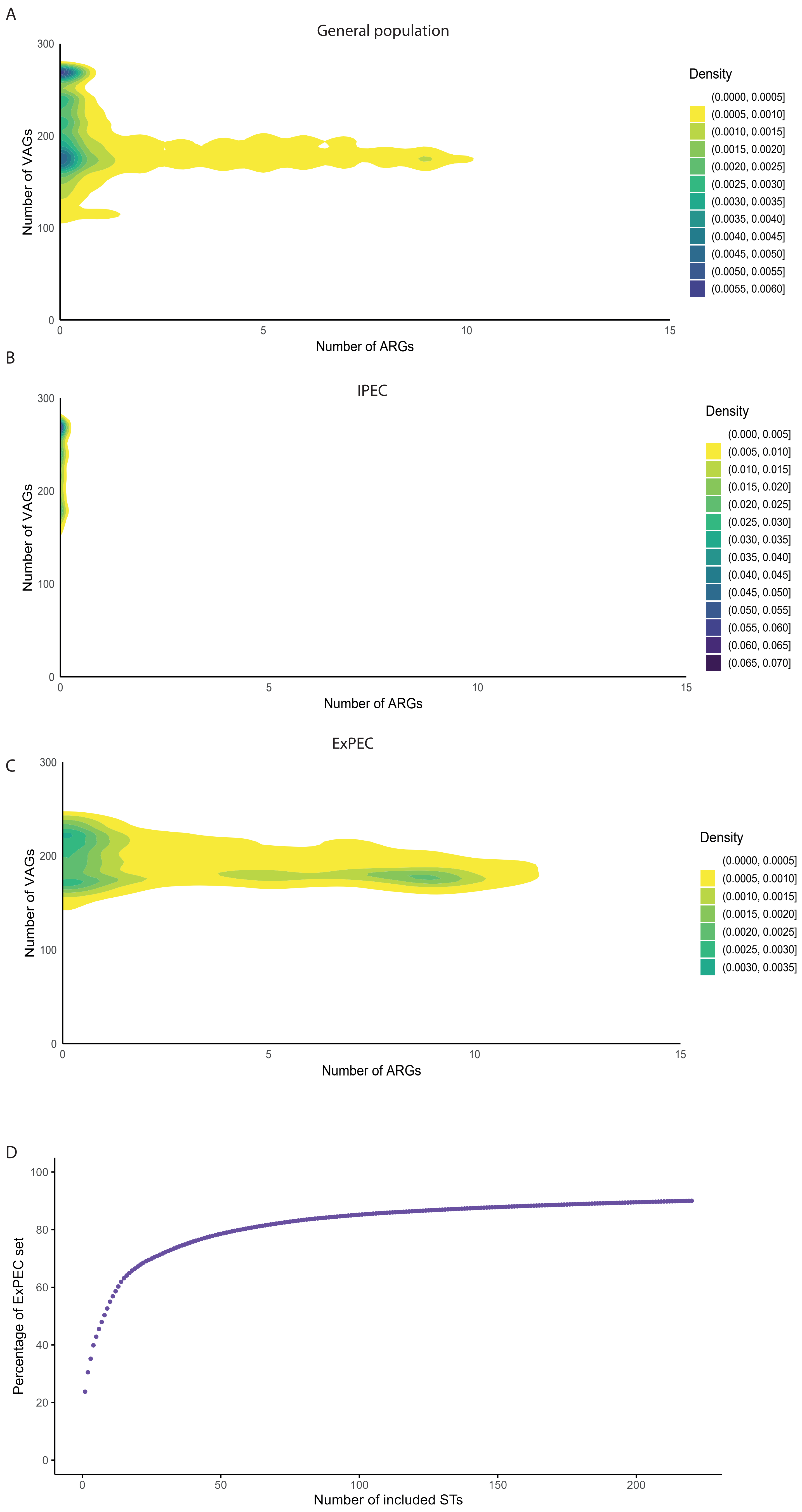
A
General population
Density
(0.0000, 0.0005]
(0.0005, 0.0010]
(0.0010, 0.0015]
(0.0015, 0.0020]
(0.0020, 0.0025]
(0.0025, 0.0030]
(0.0030, 0.0035]
(0.0035, 0.0040]
(0.0040, 0.0045]
(0.0045, 0.0050]
(0.0050, 0.0055]
(0.0055, 0.0060]
Number of VAGs
300
200
100
0
0
5
10
15
Number of ARGs
B
IPEC
Density
(0.000, 0.005]
(0.005, 0.010]
(0.010, 0.015]
(0.015, 0.020]
(0.020, 0.025]
(0.025, 0.030]
(0.030, 0.035]
(0.035, 0.040]
(0.040, 0.045]
(0.045, 0.050]
(0.050, 0.055]
(0.055, 0.060]
(0.060, 0.065]
(0.065, 0.070]
Number of VAGs
300
200
100
0
0
5
10
15
Number of ARGs
C
ExPEC
Density
(0.0000, 0.0005]
(0.0005, 0.0010]
(0.0010, 0.0015]
(0.0015, 0.0020]
(0.0020, 0.0025]
(0.0025, 0.0030]
(0.0030, 0.0035]
Number of VAGs
300
200
100
0
0
5
10
15
Number of ARGs
D
Percentage of ExPEC set
100
80
60
40
20
0
0
50
100
150
200
Number of included STs
