## Supplementary material for "From species-wide genomics to cellular diversity: characterizing multidrug-resistant extraintestinal pathogenic *Escherichia coli* ST744": Figure S2

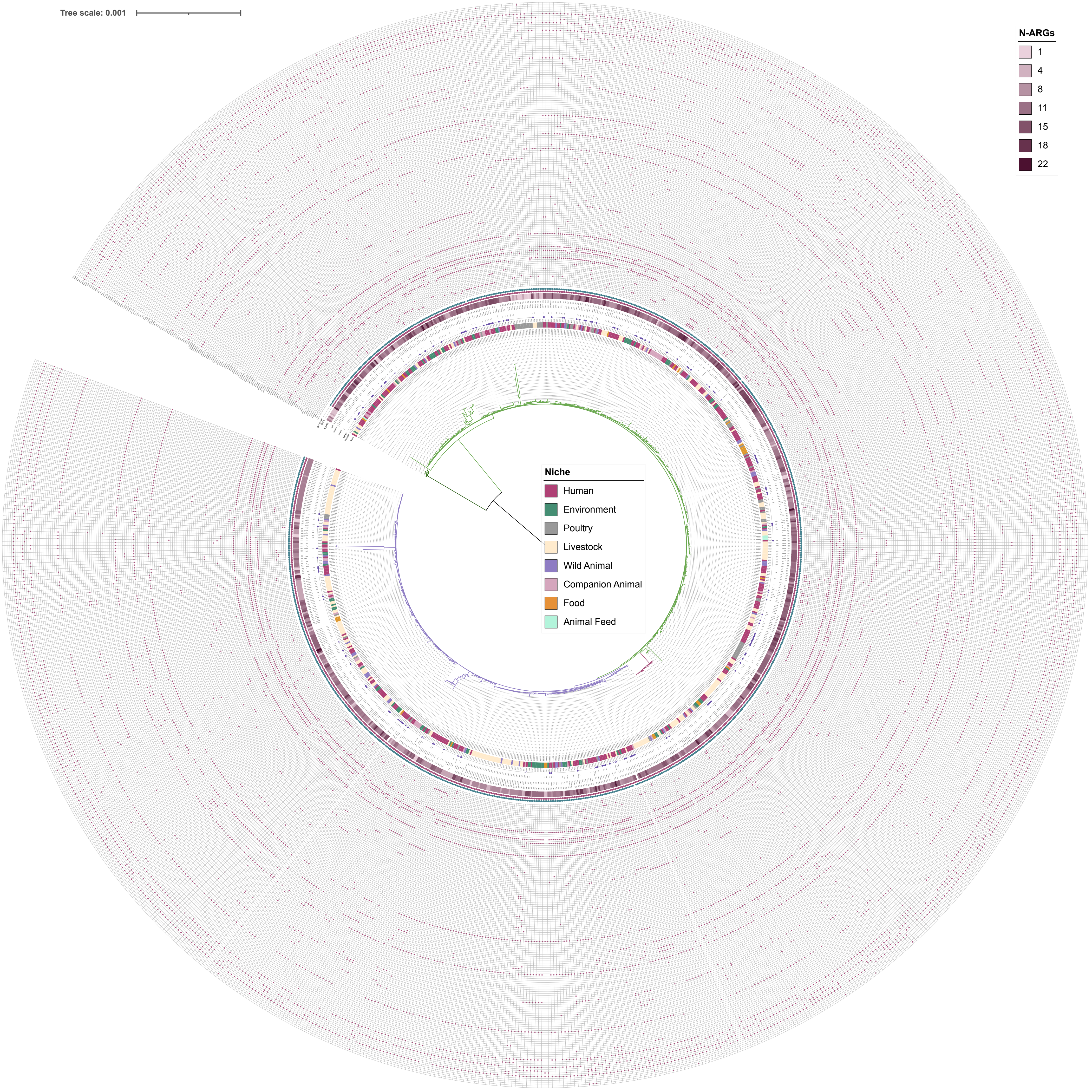
Tree scale: 0.001
N-ARGs
1
4
8
11
15
18
22
Niche
Human
Environment
Poultry
Livestock
Wild Animal
Companion Animal
Food
Animal Feed
