## Supplementary material for "From species-wide genomics to cellular diversity: characterizing multidrug-resistant extraintestinal pathogenic *Escherichia coli* ST744": Figure S3

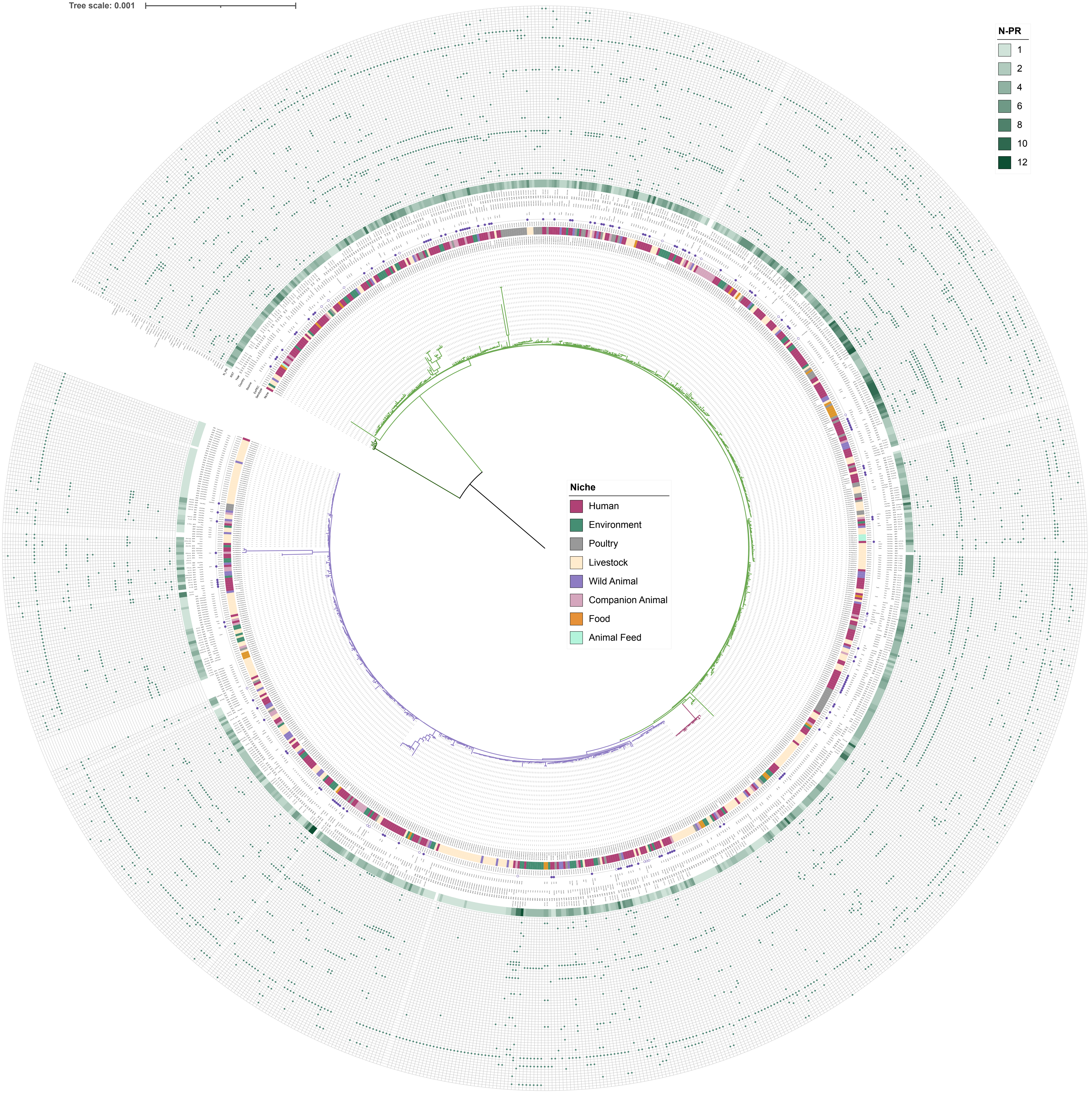

Tree scale: 0.001
N-PR
1
2
4
6
8
10
12
Niche
Human
Environment
Poultry
Livestock
Wild Animal
Companion Animal
Food
Animal Feed
