## Supplementary material for "From species-wide genomics to cellular diversity: characterizing multidrug-resistant extraintestinal pathogenic *Escherichia coli* ST744": Figure S5

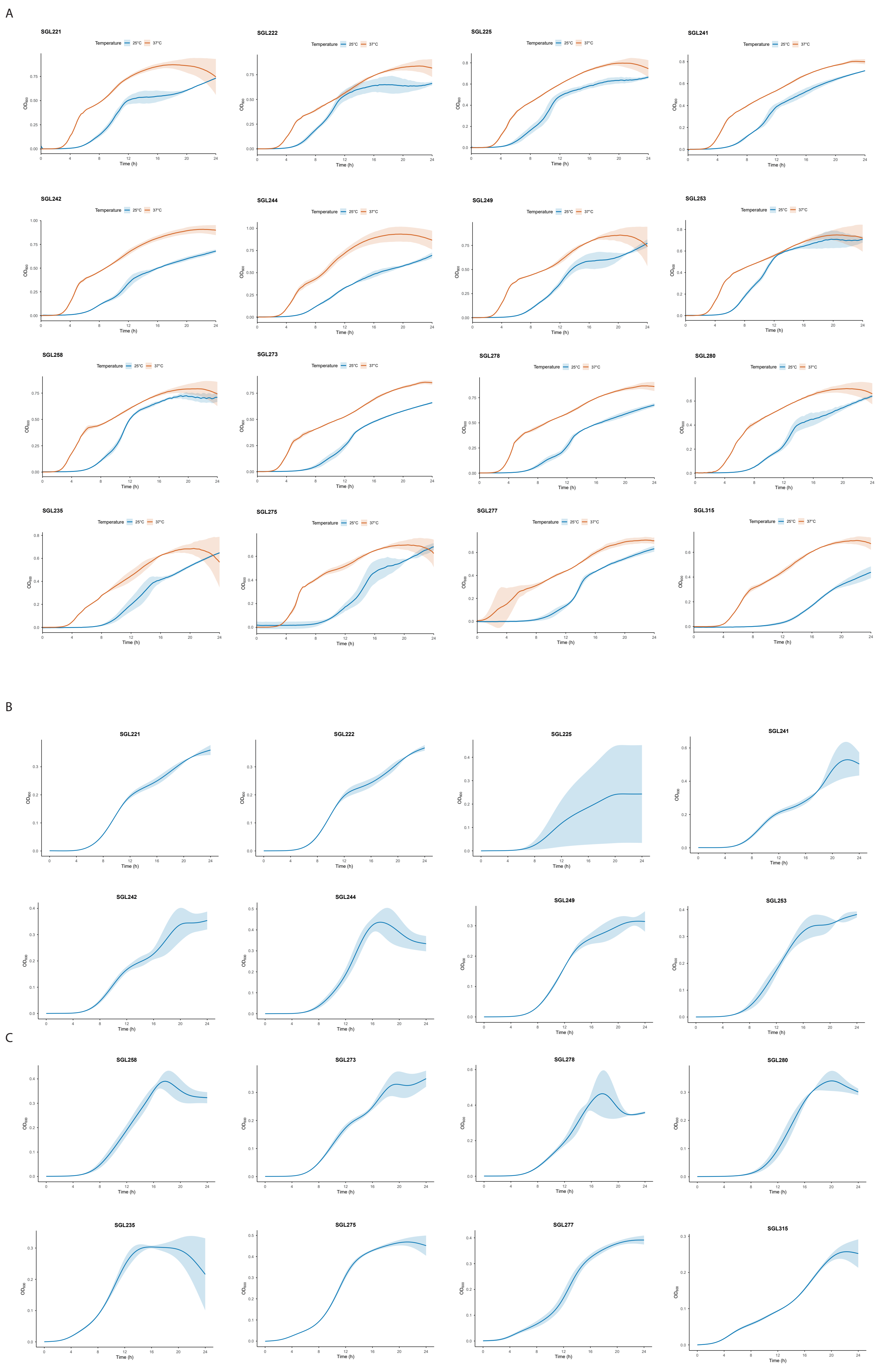
A
SGL221
SGL222
SGL225
SGL241
SGL242
SGL244
SGL249
SGL253
SGL258
SGL273
SGL278
SGL280
SGL235
SGL275
SGL277
SGL315
Temperature
25°C
37°C
OD600
Time (h)
B
SGL221
SGL222
SGL225
SGL241
SGL242
SGL244
SGL249
SGL253
C
SGL258
SGL273
SGL278
SGL280
SGL235
SGL275
SGL277
SGL315
OD600
Time (h)
