## Supplementary figures_legends for "From species-wide genomics to cellular diversity: characterizing multidrug-resistant extraintestinal pathogenic *Escherichia coli* ST744"

**Figure S1: Probability density of genomes distribution based on VAGs and ARGs counts**

A) Probability density based on whole EB collection.
B) Probability density based on IPEC genomes.
C) Probability density based on ExPEC genomes.
D) Coverage in percentage of the ExPEC collection for the 222 first most dominant ExPEC STs.

**Figure S2: Phylogenetic tree showing the presence of all detected resistance genes**

The branches are coloured according to the dominant serotype: H9:O101 (green, a distinct clade of this serotype in dark green), H10:O101 (violet) and H25:O101 (pink). The metadata in order from the most inner circle resents Niche, Serotype, ExPEC status (ExPEC, with full violet circle representing ExPEC-labelled strains, while empty violet circle represent additional clinically relevant strains without further specification), Source of isolation (Source), Country of origin (Country) and Year of isolation (Year). The number of detected ARGs (N_ARGs) is shown in a form of a heatmap and the presence of specific markers is shown as full circle/square including QRDR point mutations (pink circle), and a point mutation causing resistance to fosfomycin (*glpT*_E448K, turquoise circle) and the presence of detected ARGs (pink square).

**Figure S3: Phylogenetic tree showing the presence of all detected plasmids replicons**

The branches are coloured according to the dominant serotype: H9:O101 (green, a distinct clade of this serotype in dark green), H10:O101 (violet) and H25:O101 (pink). The metadata in order from the most inner circle resents Niche, Serotype, ExPEC status (ExPEC, with full violet circle representing ExPEC-labelled strains, while empty violet circle represent additional clinically relevant strains without further specification), Source of isolation (Source), Country of origin (Country) and Year of isolation (Year) and RST of F plasmids (RST). The number of detected plasmid replicons (N_PR) is highlighted in the form of a heatmap and the presence plasmid replicons is shown as a full square (turquoise).

**Figure S4: Map of a complete F18:A:B1 plasmid.**

The most dominant F plasmid RST F18:A:B1 detected in our set of 892 ST744 represented by a complete plasmid of 127131 bp from SGL242.

**Figure S5: Growth rates comparison for 12 selected ST744 strains and 4 comparative strains (ST69, ST155, ST354 and MG1655)**

The curves are visualized together with ±1 standard deviation across biological replicates.

A) Growth curves at 37 °C (orange) and 25 °C (blue) for the set of 12 selected ST744 strains and comparative strains (last row) of ST69 (SGL275), ST155 (SGL277), ST354 (SGL235) and MG1655 (SGL315).
B) Growth curves at 37 °C in M9 supplemented with fucose for the set of 12 selected ST744 strains and comparative strains (last row) of ST69 (SGL275), ST155 (SGL277), ST354 (SGL235) and MG1655 (SGL315).
