## Supplementary tables_legends for "From species-wide genomics to cellular diversity: characterizing multidrug-resistant extraintestinal pathogenic *Escherichia coli* ST744"

**Table S1: Metadata of 892 *E. coli* ST744 strains and results from ResFinder, PlasmidFinder, VFDB, and pMLST.**

The metadata include curated information on our 32 ST744 strains and strains obtained from EnteroBase (identified by the ESC_ prefix in the Strain ID). The table provides counts of detected antimicrobial resistance genes (ARGs), plasmid replicons, and virulence-associated genes (VAGs; N_ARGs, N_PR, and N_VAGs), as well as an overview of ARGs and resistance-associated point mutations detected by ResFinder (blue background), F-plasmid replicon sequence types (RSTs) determined by pMLST, plasmid replicons detected by PlasmidFinder (green background), and VAGs detected by VFDB.

**Table S2: Gene presence–absence table generated by Roary for 892 ST744 genomes.**

The presence (strain name and corresponding contig) or absence (empty field) of each gene in the ST744 pangenome is shown for each strain (blue background).

**Table S3: SNP matrix.**

SNP matrix for 892 ST744 strains, with clonal pairs (up to 10 SNPs apart) highlighted in red.

**Table S4: Selected clonal pairs.**

The table highlights the One Health- and epidemiological relevance of ST744 by listing pairs or groups of three strains from the same (or neighbouring) country and, in most cases, the same year. For groups of three strains, the strain highlighted in pink is used as the reference for SNP comparisons with the other two strains (forming two clonal pairs).

**Table S5: Genome-wide association analysis for the H10 serotype.**

Genes strongly associated with H10 are listed on a separate sheet.

**Table S6: Genome-wide association analysis for the H9 serotype.**

Genes strongly associated with H9 are listed on a separate sheet.

**Table S7: Prevalence of selected genomic markers.**

Prevalence is shown for the set of 892 ST744 genomes.

**Table S8: Prevalence of heavy-metal resistance genes.**

The table shows the presence of genes associated with resistance to arsenic and antimony, mercury, tellurite, silver, copper, nickel, and cobalt across 892 ST744 genomes. Genomes with no detected genes are not included.

**Table S9: Minimum inhibitory concentrations for 32 ST744 isolates.**

The table includes minimum inhibitory concentrations (MICs) for 24 antibiotics or antibiotic combinations. Where applicable, thresholds for resistant (pink background), susceptible (green background), and intermediate (blue background) phenotypes are highlighted.

**Table S10: Prevalence of insertion sequences in ST744 compared with the general *E. coli* population.**

The *Comparison* sheet shows the prevalence (%) of the listed insertion sequences (ISs) in ST744 and compares ISs with a prevalence above 90% with their prevalence in the general *E. coli* population (data from Table S20). ISs with a prevalence difference greater than 50 percentage points are highlighted in yellow. The *Prevalence* sheet shows the prevalence of the listed ISs in ST744 genomes.

**Table S11: Prevalence of cold-shock genes in ExPEC STs.**

The *Summary* sheet shows the prevalence of five cold-shock genes associated with ST744 in other typical ExPEC STs, while the subsequent sheets provide the corresponding data for each included ST individually.

**Table S12: Prevalence of regions of difference identified in the accessory genomes of two reference strains.**

The prevalence of genes from regions of difference identified by comparing two reference strains (SGL242 and SGL280, each presented on a separate sheet) with MG1655 is shown across 892 ST744 genomes.

**Table S13: Prevalence of ExPEC-associated genes and AW-ROD genes.**

Genes from Table S11 that are associated with ExPEC (a prevalence difference of at least 20 percentage points compared with the ND group) or belong to region of difference AW.

**Table S14: ExPEC-associated genes identified by pangenome screening.**

ExPEC-associated genes identified by screening 16,226 accessory genes across 892 ST744 genomes.

**Table S15: ST744 core genes uncommon in *E. coli*.**

ST744 core genes with low prevalence (below 10%) in the general *E. coli* population.

**Table S16: Genome-wide association analysis for ExPEC within ST744.**

The analysis is based on a subset of 500 genomes from the full collection for which clinical status could be determined.

**Table S17: Prevalence of selected genomic markers in strains subjected to growth-curve experiments.**

Prevalence of cold-shock genes and AW-ROD genes in 12 selected ST744 isolates and four comparison strains (three different STs and MG1655).

**Table S18: Mean features extracted from microscopy data.**

Mean features extracted from microscopy data using BactoScoop for 32 ST744 isolates and MG1655.

**Table S19: Z-scores of 14 selected features for 32 ST744 isolates and MG1655.**

Z-scores of 14 selected features extracted from microscopy data for 32 ST744 isolates and MG1655.

**Table S20: Z-scores of 14 selected features for 32 ST744 isolates.**

Z-scores of 14 selected features extracted from microscopy data for 32 ST744 isolates.

**Table S21: Metadata of ST744 from EnteroBase.**

Original, non-curated metadata of ST744 from EnteroBase.

**Table S22: Prevalence of insertion sequences in *E. coli*.**

Prevalence of insertion sequences in the general *E. coli* population, based on a dataset of 1.933 complete *E. coli* genomes from NCBI.
